# Single particle cryo-EM structure of the outer hair cell motor protein prestin

**DOI:** 10.1101/2021.08.03.454998

**Authors:** Carmen Butan, Qiang Song, Jun-Ping Bai, Winston J. T. Tan, Dhasakumar Navaratnam, Joseph Santos-Sacchi

## Abstract

The mammalian outer hair cell (OHC) protein prestin (Slc26a5), a member of the solute carrier 26 (Slc26) family of membrane proteins, differs from other members of the family owing to its unique piezoelectric-like property that drives OHC electromotility. OHCs require prestin for cochlear amplification, a process that enhances mammalian hearing. Despite substantial biophysical characterization, the mechanistic basis for the prestin’s electro-mechanical behavior is not fully understood. To gain insight into such behavior, we have used cryo-electron microscopy at subnanometer resolution (overall resolution of 4.0 Å) to investigate the three-dimensional structure of prestin from *gerbil* (*Meriones unguiculatus*). Our studies show that prestin dimerizes with a 3D architecture strikingly similar to the dimeric conformation observed in the Slc26a9 anion transporter in an inside open/intermediate state, which we infer, based on patch-clamp recordings, to reflect the contracted state of prestin. The structure shows two well-separated transmembrane (TM) subunits and two cytoplasmic sulfate transporter and anti-sigma factor antagonist (STAS) domains forming a swapped dimer. The dimerization interface is defined by interactions between the domain-swapped STAS dimer and the transmembrane domains of the opposing half unit, further strengthened by an antiparallel beta-strand at its N terminus. The structure also shows that each one of its two transmembrane subunits consists of 14 transmembrane segments organized in two inverted 7-segment repeats with a topology that was first observed in the structure of bacterial symporter UraA (Lu F, et al., Nature 472, 2011). Finally, the solved anion binding site structural features of prestin are quite similar to that of SLC26a9 and other family members. Despite this similarity, we find that SLC26a9 lacks the characteristic displacement currents (or **N**on**L**inear **C**apacitance(NLC)) found with prestin, and we show that mutation of prestin’s Cl^-^ binding site removes salicylate competition with anions in the face of normal **NLC**, thus refuting the yet accepted extrinsic voltage sensor hypothesis and any associated transport-like requirements for voltage-driven electromotility.

## Introduction

The outer hair cell (OHC) molecular motor prestin (Slc26a5) belongs to a diverse family of transporters that includes Slc26, Slc4 and Slc23 (Chang and Geertsma, 2017). Unlike other members of these families, and unique to membrane proteins, prestin functions as a voltage-driven motor with rapid kinetics, likely providing cycle-by-cycle amplification of sound within the mammalian organ of Corti (Santos-Sacchi et al., 2006; Dallos et al., 2008). However, cycle-by-cycle amplification at frequencies higher than 50 kHz, where mammals such as bats and whales can hear, may be limited by the low pass nature of prestin’s voltage sensor charge movement, which is a power function of frequency that is 40 dB down (1 %) in magnitude at 77 kHz (Santos-Sacchi and Tan, 2018, 2019, 2020). The underlying basis of prestin’s electromechanical capabilities resides in its unique piezoelectric-like property that drives OHC electromotility (Iwasa, 1993; Gale and Ashmore, 1994; Kakehata and Santos-Sacchi, 1995; Ludwig et al., 2001; Santos-Sacchi et al., 2001). For members of this diverse family, known structures are dimers with each protomer showing a common 7+7 inverted repeat topology containing a core and gate domain; these proteins function variably as ion channels and transporters with a range of substrates (Chang and Geertsma, 2017; Yu et al., 2017; Chang et al., 2019). Within the Slc26 family, prestin and pendrin (Slc26a4) are unique in showing voltage sensitivity with signature nonlinear capacitance (NLC) or equivalently, displacement currents/gating charge movements (Ashmore, 1990; Santos-Sacchi, 1991; Kuwabara et al., 2018); while pendrin lacks intrinsic electromechanical behavior (Tang et al., 2011), prestin is a minimal transporter (Ashmore, 2005; Bai et al., 2009; Schanzler and Fahlke, 2012). With more structural information there have been competing visions of transporter mechanisms (elevator vs. rocker) (Drew and Boudker, 2016; Ficici et al., 2017), although how these fit with prestin’s electromechanical behavior remains speculative at best. To be sure, the lack of structural information for full-length prestin has precluded an understanding of its unique molecular motor function. In this study, we have used single-particle cryo-electron microscopy to determine the structure of prestin from gerbil (*Meriones unguiculatus*) at sub-nanometer resolution that confirms Oliver’s initial modeling efforts (Gorbunov et al., 2014) and, remarkably, bears high concordance with the recently determined cryo-EM structures of Slc26a9 (Walter et al., 2019; Chi et al., 2020). Prestin forms a dimer and the cryo-EM density map has allowed us to build a nearly complete model of the protein. In combination with electrophysiological data, our structural results suggest that the inward cytosol facing conformation is that of prestin in the contracted state. Furthermore, mutations within prestin’s now structurally confirmed anion binding site show that the extrinsic voltage sensor hypothesis (Oliver et al., 2001) is likely incorrect (Santos-Sacchi et al., 2017), with the wider implication that a transporter-like mechanism driving electromotility is unlikely.

## Results

### The general architecture of full-length Prestin

We used single-particle cryo-electron microscopy to obtain the structure of detergent-solubilized prestin from gerbil (*Meriones unguiculatus*) (**Figure 1**). The protein, extracted digitonin, and purified in the presence of GDN (glyco-diosgenin), appears to be a homogeneous oligomer when assessed with size-exclusion chromatography (**Figure 1, Supplement 1**).

**Figure 1.**
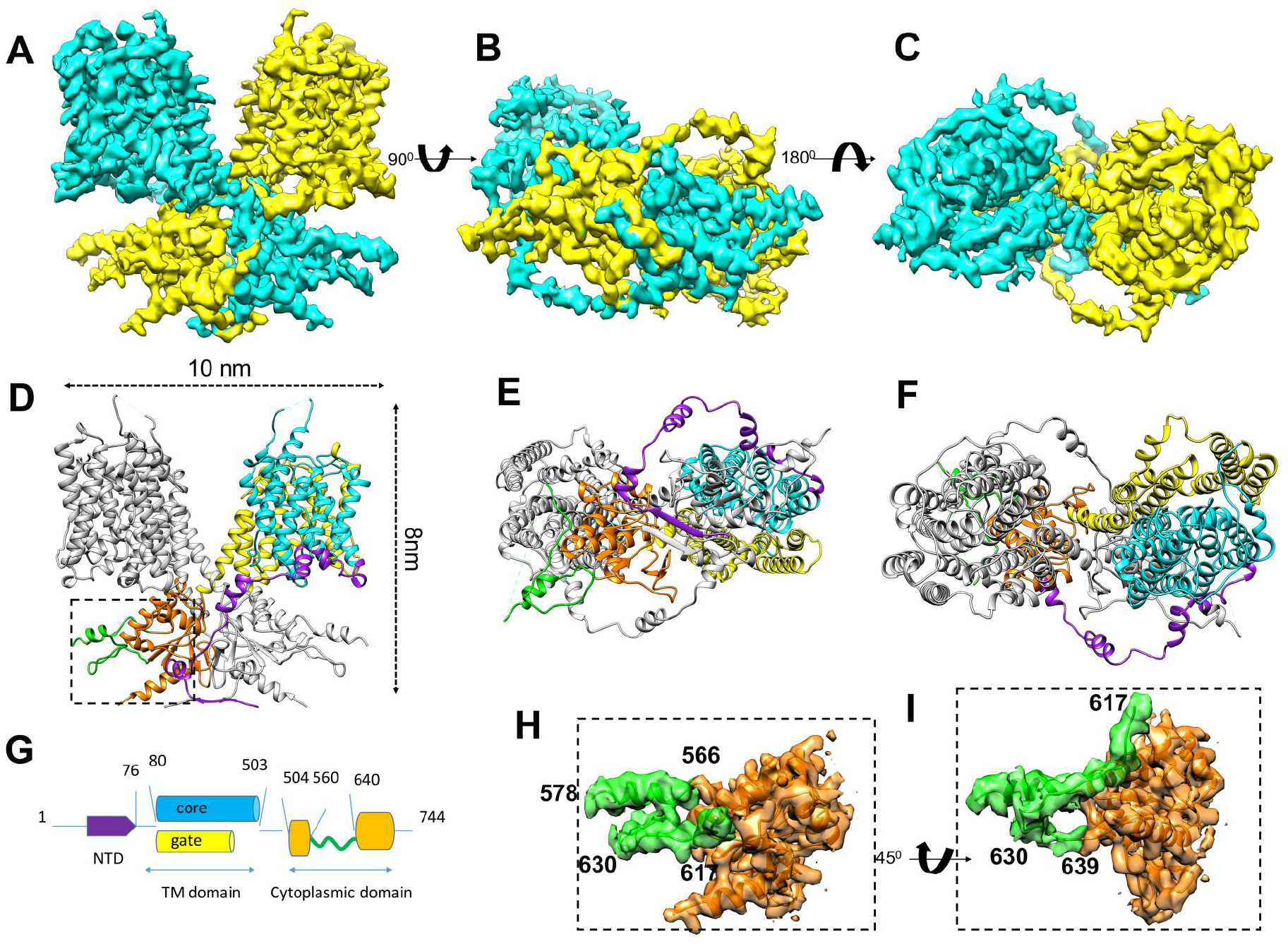
The cryo-EM structure of prestin from gerbil. (**A-C**) Three views: side view (**A**), cytosolic view (**B**) and extracellular view (**C**) of the cryo-EM structure of the prestin dimer. The prestin structure is colored by subunit in cyan and yellow respectively. (**D-F**) Atomic model based on the cryo-EM density, shown in the same orientations as the density maps in the panels (**A-C**). The different domain structures of prestin are colored using the same color scheme as shown in the schematic representation of the prestin sequence (**G**). (**H**) and (**I**) Two close-up views, rotated ∼45° with respect to each other, showing a density segment of the IVS loop overlaid with the model (shown in green) within the STAS domain structure (shown in orange). Panels (**H, I**) show details of the interface boxed in panel (**D**).

The cryo-EM images obtained from plunge-frozen specimens of solubilized prestin (**Figure 1, Supplement 2A**) revealed clear density for the transmembrane helices (TM), the cytoplasmic domains, and the micelle belt around the protein (**Figure 1, Supplement 2B**). A density map (**Figure 1A-C**) was obtained at an overall resolution of 4.0 Å according to the gold-standard Fourier shell correlation (FSC) from 122,754 particles using C2 symmetry (**Figure 1, Supplement 2C**,**D**). The density map displayed clear secondary structural elements and densities visible for many of the bulky side chains (**Figure 1, Supplement 2E**,**F**). An analysis of the local-resolution of the EM map shows that the interior of the structure is better resolved than the periphery (**Figure 1, Supplement 2C**,**D**), with the lowest resolution observed in the extracellular loops, the cytosolic IVS loop (variable or intervening sequence) and the cytosolic C-terminal domain. This map has allowed us to unambiguously build and refine a nearly complete model of full-length prestin (**Figure 1D**,**E**,**F**). The cryo-EM derived structure shows that prestin oligomerizes as a dimer, with overall dimensions of ∼10 nm in diameter and ∼8 nm in height. Each protomer comprises 14 transmembrane helices (named TM1-TM14), a C-terminal cytosolic STAS domain, and a short cytosolic N-terminal region. The 14 transmembrane segments exhibit the same inverted 7-segment repeat organization as first observed in the crystal structure of the bacterial symporter UraA (Lu F, et al., Nature 472, 2011). Seven of the transmembrane segments (TM 1-4, TM 8-11), are referred to as the “core” domain and the remaining seven helices (TM 5-7, TM 12-14) form the “gate” domain of the structure. The largest interaction between prestin subunits is made by the contacts between the STAS domain of one subunit (depicted by the grey ribbon in the side view of prestin structure as seen in **Figure 1D**) and the transmembrane domain of the opposing half subunit (represented by the yellow and cyan ribbons in the side view of prestin structure as seen in **Figure 1D**). The majority of this interface is formed by residues located at the tips of TM5, TM8 and TM12-14 helices, which are exposed to the cytosol, and the residues located on the first 3 helices of the opposing STAS domain. Close contacts between residues located on the anti-parallel beta-strands at the N-terminal domain of each prestin subunit also contribute to the inter-subunit interactions (**Figure 1E**).

### Comparison of the cryo-EM structure of prestin with the cryo-EM structure of Slc26a9

The cryo-EM structure of prestin closely resembles the previously reported cryo-EM structure of Slc26a9, which is a representative Slc26 family member (Walter et al., 2019; Chi et al., 2020). In particular, the dimer of Slc26a9 from the mouse in “intermediate” conformation (PDB ID, 6RTF, pair-wise C_α_ RMSDs: 2.985 Å overall) can be fitted into the cryo-EM density map of prestin from gerbil reasonably well. The previously observed Slc26a9 “intermediate” conformation (Walter et al., 2019) may represent a more closed conformation when compared to the “inward-open” conformation of Slc26a9 (Walter et al., 2019). Our prestin structure superimposes onto the “inward-open” conformation of Slc26a9 from mouse (PDB ID, 6RTC), with pair-wise C_α_ RMSDs of 4.25 Å overall. Of note, the two previously resolved “inward-open” conformations of human SLC26A9 (PDB ID, 7CH1, determined at a resolution of 2.6 Å) and mouse Slc26a9 (PDB ID, 6RTC, determined at a resolution of 3.96 Å resolution) are essentially identical, with only very subtle changes.

When the prestin monomer was superimposed onto the Slc26a9 monomer in “intermediate” and “inward-open” conformations, through their respective STAS domains (**Figure 2, Supplement 1 A**,**B**,**C**), the first large displacements with the Slc26a9 structure are observed at the linker connection between the transmembrane and cytosolic domains, indicated by an arrow. The displacements extend into the entire transmembrane domains (**Figure 2, Supplement 1 A**,**B**,**C**). An intriguing result becomes apparent when superimposing only the separated transmembrane domain of prestin onto the transmembrane domains of the previously solved Slc26a9 cryo-EM structures (**Figure 2, Supplement 1 D**,**E**,**F**) through the C_α_ atoms of residues within TM13 and TM14 helices. We observe differences in the orientation of TM8 (from the core domain) with respect to the TM13 and TM14 (from the gate domain) (**Figure 2 E**,**F**,**G; Figure 2, Supplement 2 A**,**B**,**C**). In prestin, TM8 moves closer to the TM14, whereas in the Slc26a9 “inward-open” conformation, TM8 splays apart from TM14. This causes the gap between the ends of TM13, TM14 (from the gate domain), and the end of TM8 (from the core domain) to be the larger in the “Inward-open” conformation of Slc26a9 and the smaller in the prestin conformation resolved by our cryo-EM structure (**Figure 2 E**,**F**,**G; Figure 2, Supplement 2 A**,**B**,**C**). We speculate that our cryo-EM prestin conformation, showing a narrower opening to the cytosol, would explain why this protein shows reduced ability to transport anions in comparison to Slc26a9. The RMSD values when superimposing the transmembrane domain C_α_ atoms of prestin residues within TM13 and TM14 are 4.02 Å (Slc26a9, PDB ID, 7CHR1) and 3.9 Å (Slc26a9, PDB ID, 6RTF), respectively.

**Figure 2.**
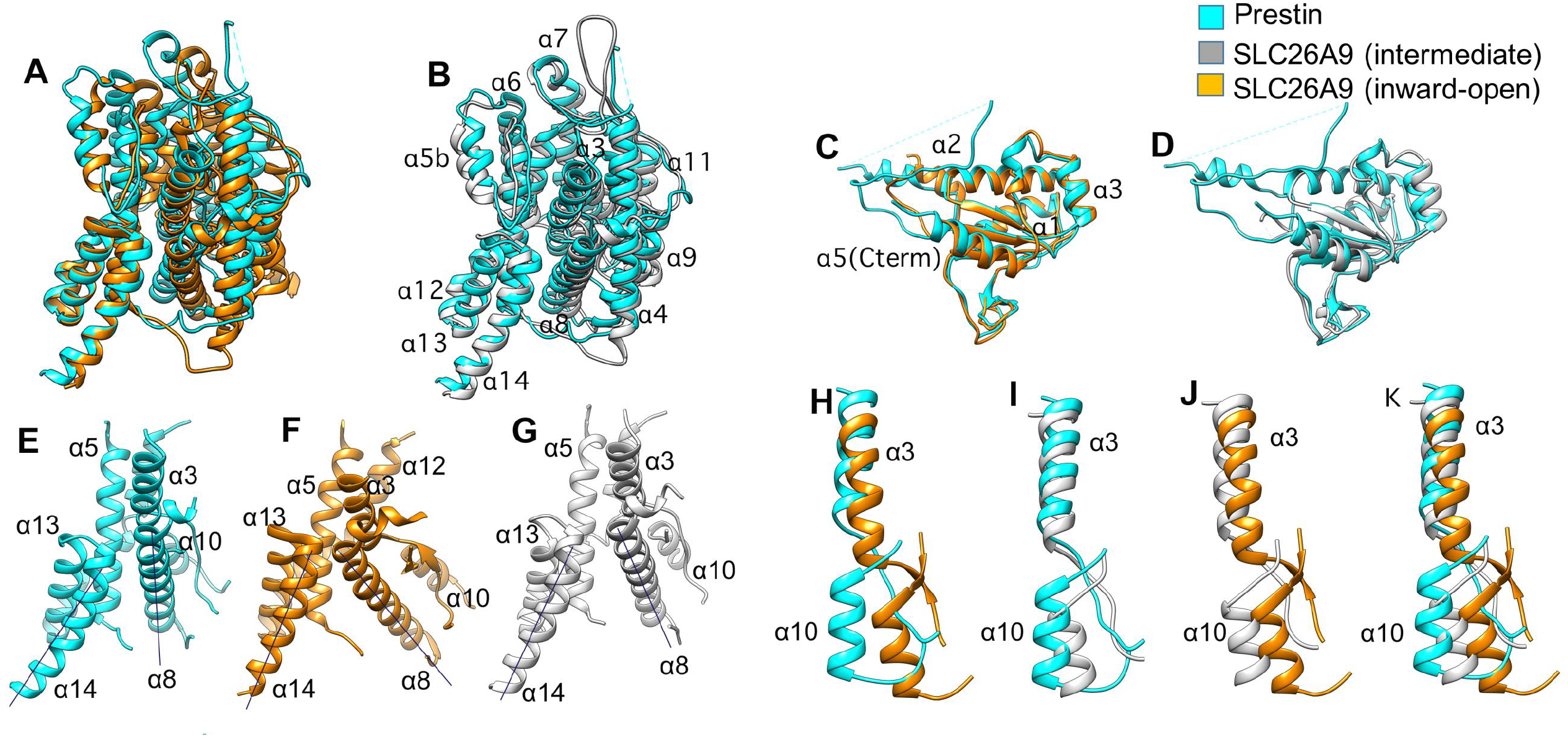
Comparison of the cryo-EM prestin structure to the cryo-EM Slc26a9 structure (**A**) Structural comparison of the transmembrane domain from the prestin structure (in cyan) with the transmembrane domain of the “inward-open” conformation of the Slc26a9 structure (PDB ID, 7CH1, in orange). The structures have been superimposed through the TM13 and TM14 helices from the “gate” domain. (**B**) Structural comparison of the transmembrane domain from the prestin structure (in cyan) with the transmembrane domain of the “intermediate” conformation from the Slc26a9 structure (PDB ID, 6RTF, in grey) (superimposed through the TM13 and TM14 from the “gate” domain). (**C, D**) Structural comparison of the STAS domain from prestin (in cyan) with the STAS domains from the two conformations of Slc26a9 (in orange and in cyan). (**E, F, G**) Side by side comparisons of the orientations of the TM8 helix with respect to TM13 and TM14 helices from the prestin structure (in cyan) and Slc26a9 structures (in orange and in grey). For clarity, line segments indicate the orientations of TM8, TM13 and TM14 helices. (**H, I, J, K**) Overlays illustrating the offset between the pair of helices (TM3, TM10) from the prestin structure (in cyan) and the (TM3, TM10) helices from the “inward-open” conformation of Slc26a9 (PDB ID, 7CH1, in orange) and “the intermediate” conformation of Slc26a9 (PDB ID, 6RTF, in grey).

The C-terminal STAS domain of prestin is similar to the previously determined X-ray crystal structure (PDB ID, 3LLO), lacking the unstructured loop (Pasqualetto et al., 2010). Thus, we see an identical core of five beta-sheets surrounded by 5 alpha-helices. The RMSD value when superimposing prestin’s C-terminal domain solved by cryo-EM with prestin’s C-terminal domain solved by X-ray crystallography (PDB ID, 3LLO) is 0.853 Å over 105 C_alpha_ atoms. The prestin’s STAS domain and the STAS domains of the two Slc26a9 conformations align with an RMSD value of 0.943 Å (PDB ID, 7CHR1) and of 1.038 Å respectively (PDB ID, 6RTF), revealing essentially identical structures (**Figure 2C**,**D; Figure 2 Supplement 1 G**,**H**,**I**).

As opposed to the previously reported SLC26a9 cryo-EM structures (Walter et al., 2019; Chi et al., 2020), the cryo-EM density for the IVS loop (variable or intervening sequence) resolved in prestin shows a more ordered, interpretable density (**Figure 1, H**,**I**). As shown in **Figure 1, H**,**I**, the IVS sequence (residues 617 to 630) abuts on the tip of helix 2 (residues G668, D669) and the loop connecting helix3 with helix 4 (residues E694, N695) from the STAS domain.

### Functional implications of prestin’s structure

#### Prestin is in the contracted state

We infer that the structure of prestin that we obtained is in the contracted state. Membrane depolarization from a negative resting voltage shifts prestin from an expanded to contracted state, evoking OHC shortening (Ashmore, 1990; Santos-Sacchi, 1991). Furthermore, increases in intracellular Cl^-^ ion concentration cause a hyperpolarizing shift in V_h_, the voltage where, based on 2-state models, prestin is equally distributed between compact and expanded states (Oliver et al., 2001; Rybalchenko and Santos-Sacchi, 2003b; Rybalchenko and Santos-Sacchi, 2003a). The cell line from which we purified prestin (Bian et al., 2010) displays V_h_ values near -110 mV in the presence of 140 mM Cl^-^ (**Figure 3A**). Since the voltage across detergent micelles is effectively 0 mV, prestin must be in a contracted state.

**Figure 3.**
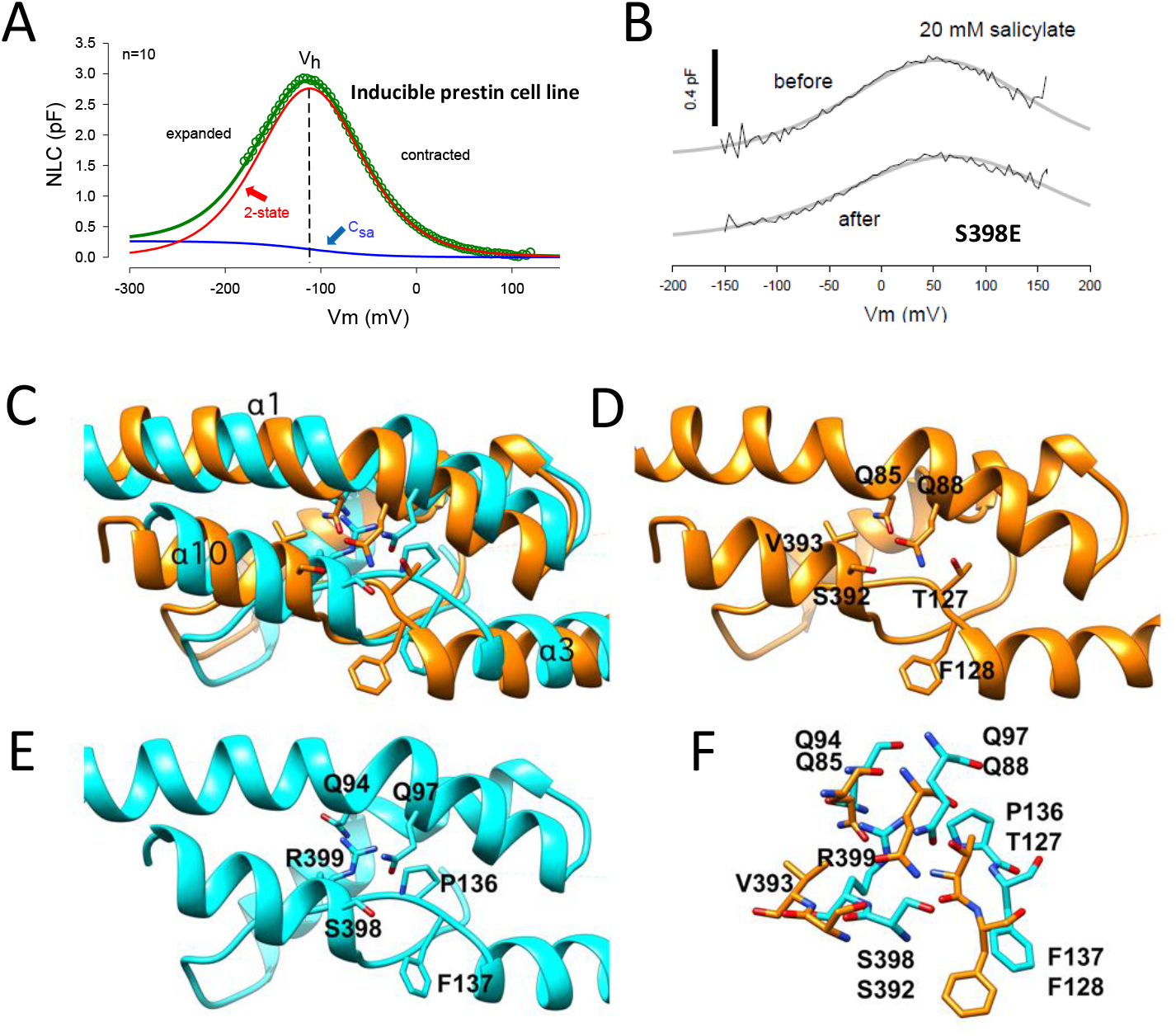
**(A)** Prestin predominantly resides in the contracted state at 0 mV. Average NLC (n=10), 48 hours after induction of our inducible prestin cell line. Recordings were done in the presence of 140 mM intracellular Cl^-^. NLC fitted (eq. 1, green line) parameters were Q_max_ = 0.40; z = 0.70; V_h_ =-112 mV; ΔC_sa_ = 0.26 pF. C_sa_ component of fit (blue line), with ΔC_sa_ describing the maximal change in linear capacitance at very negative potentials; 2-state component of fit (red line). NLC peaks at V_h_. **(B)** The S398E mutation in prestin preserves NLC after application of 20mM salicylate. Controls shows full block of NLC. The average unitary gating charge (z) in these mutants (0.69e +/-0.03 SEM, n =7) was similar to that of CHO cells expressing prestin-YFP (0.73e +/-0.14 SEM, n=10, P>0.05). **(C)** Overlay of the Slc26a9’s TM segments (TM1, TM3, TM10, in orange) with the equivalent prestin’s TM segments (TM1, TM3, TM10 in cyan) showing residues which are important for the Cl-binding. **(D)** Mapping of Q85, Q88, S392, T127, F128 residues which have been demonstrated to be involved in anion coordination on the Slc26a9 structure. **(E)** Mapping of the equivalent residues important for substrate binding Q94, Q97, S398, R399, P136, F137 on the prestin structure. **(F)** Superposition of the residues important for substrate binding in prestin (in cyan) and Slc26a9 (orange).

#### Analysis of mutations in the chloride binding site

Despite purification in high chloride, we were unable to resolve a density corresponding to a chloride ion within prestin. A similar observation was made with all three structures of Slc26a9 (Walter et al., 2019; Chi et al., 2020), and in prestin’s case may be due to its relatively poor Cl^-^ binding affinity (Oliver et al., 2001; Song and Santos-Sacchi, 2010). Nevertheless, prestin presents the canonical anion binding site features identified in other structurally solved family members where substrates are resolved. The binding site is between TM3 and TM10 and the beta-sheets preceding these. Many of the residues important for coordinating substrate binding in those proteins, including S398, F137, and R399, are located in similar positions in prestin (**Figure 3 C-F**). Furthermore, with the exception of T127 in Slc26a9 (which is a proline P136 in prestin) other residues important for coordinating water molecules for substrate interactions are also conserved in prestin (Q94, Q97, F101). Although it has been speculated that Cl^-^ acts as an extrinsic voltage sensor (Oliver et al., 2001), this speculation has never been confirmed; instead, anions have been shown to foster allosteric-like modulation of prestin activity and kinetics, with anion substitute valence showing no correlation with the magnitude of prestin’s unitary charge (Rybalchenko and Santos-Sacchi, 2003a; Rybalchenko and Santos-Sacchi, 2003b, 2008; Song and Santos-Sacchi, 2010, 2013; Santos-Sacchi and Song, 2016). It is well known that salicylate blocks NLC and electromotility (Tunstall et al., 1995; Kakehata and Santos-Sacchi, 1996), possibly by competitively displacing Cl^-^ (Oliver et al., 2001). Here we show that mutation of S398 in the structurally identified anion binding pocket of prestin to a negatively charged glutamate residue results in a protein that is insensitive to salicylate yet retains normal NLC (**Figure 3B**). This observation is further evidence against Cl^-^ acting as an extrinsic voltage sensor. We see similar effects with R399E. Indeed, recently, Oliver (Gorbunov et al., 2018) reported on S396E which shows insensitivity to salicylate as proof refuting his original extrinsic voltage sensor hypothesis. They also previously observed salicylate insensitivity with R399S (Gorbunov et al., 2014).Another mutation within the anion binding pocket of prestin, F127 to alanine, resulted in a loss of NLC or a far-right shift in its voltage sensitivity that made its detection impossible (Bai et al., 2017); membrane insertion was confirmed by detectable SCN^-^ currents. We additionally mutated three residues in prestin that are implicated in Cl^-^ binding in Slc26a9, substituting alanine residues for Q94 and Q97, and substituting a threonine residue for P136 that corresponds to T127 in Slc26a9. All these mutants have normal unitary gating charge (*z*) (Q94 0.66 +/-0.06 n=7, Q97 0.65 +/-0.04 n=5, P136 0.64 +/-0.05 n=9). These mutations of prestin’s Cl^-^ binding site, besides refuting the extrinsic voltage sensor hypothesis, also make implausible any associated transport-like requirements for voltage-driven electromotility.

#### Slc26a9 is not electromechanical

The marked structural similarity between prestin and Slc26a9 may provide insight into prestin’s electromechanical behavior. In prestin, several charged residues have been shown to affect the size of the unitary gating charge and thus contribute to voltage sensing (Bai et al., 2009). Of those twelve residues, nine are conserved in Slc26a9. We transiently expressed Slc26a9 in CHO cells but were unable to find NLC or gating currents in contrast to transiently transfected CHO cells expressing prestin (**Figure 4 A**,**B *top panels***). Slc26a9 surface expression in transfected cells was successful as demonstrated by enhanced currents in the presence of extracellular SCN^-^ (**Figure 4, Supplement 1 A**,**B**) and visualization of fluorescence on the surface of these cells expressing Slc26a9 with YFP tagged to its C-terminus (**Figure 4A, insert**). Thus, despite marked structural similarities to prestin, Slc26a9 does not mimic prestin’s electromechanical behavior.

**Figure 4.**
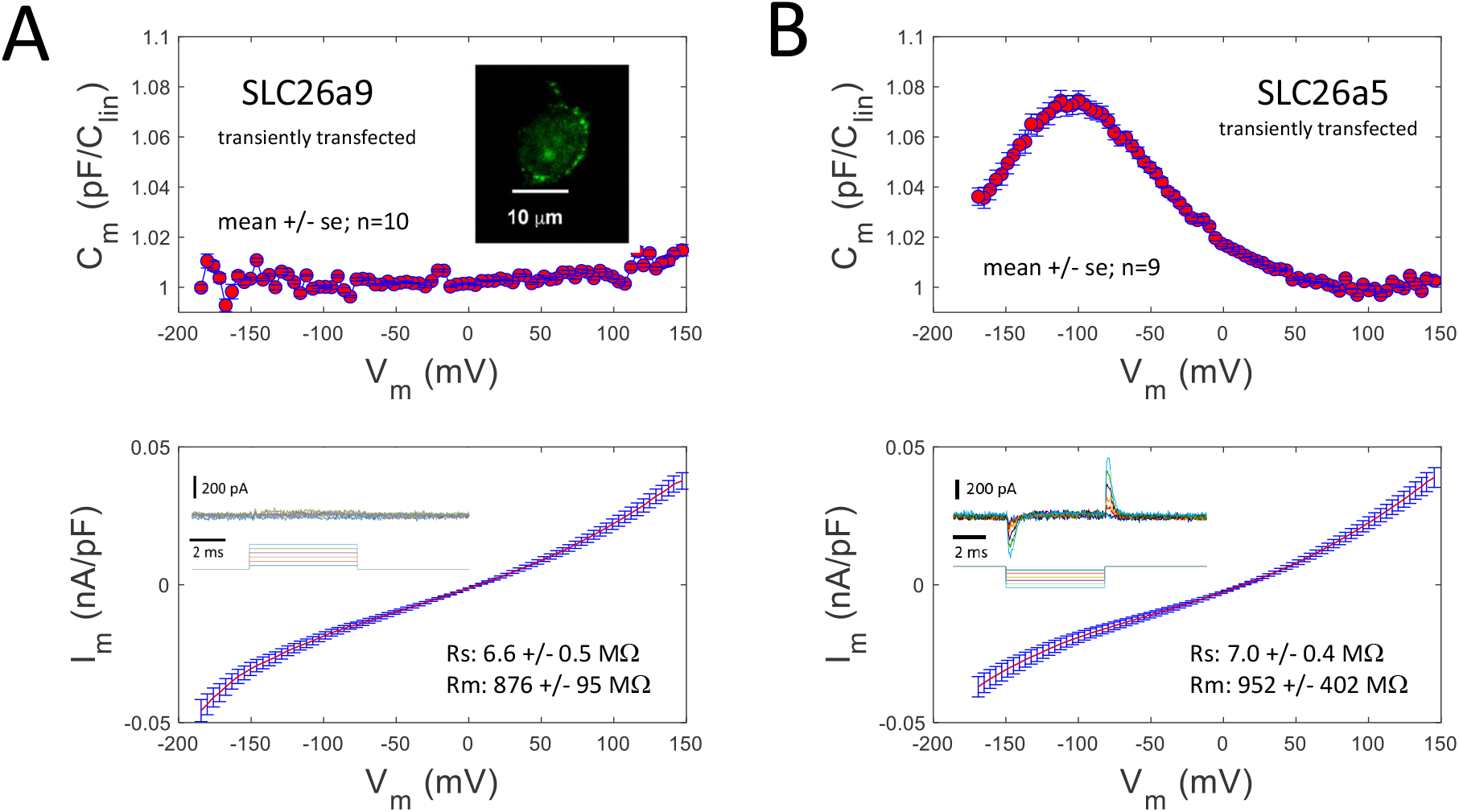
(**A**) Top *panel*: Capacitance of CHO cells transiently infected with Slc26a9, scaled to linear capacitance near -100 mV. Measured with dual-sine voltage superimposed on a voltage ramp from -175 to + 150 mV. A very slight increase in C_m_ occurs at +150 mV possibly indicating an extremely right-shifted NLC. *Insert:* Confocal image of CHO cell transfected with Slc26a9-YFP that is expressed on the membrane. The scale bar is 10 microns. *Bottom panel*: Ramp induced currents simultaneously measured with membrane capacitance, also scaled to linear capacitance. Insert: top traces show very small nonlinear currents extracted with P/-5 protocol, subtraction holding potential set to -50 mV. Voltage protocol shown below traces. (**B**) *Top panel*: Capacitance of CHO cells transiently infected with Prestin, scaled to linear capacitance near +100 mV. Measured with dual-sine voltage superimposed on a voltage ramp from -175 to + 150 mV. A prominent increase in C_m_ occurs at -110 mV, typical of prestin NLC. *Bottom panel*: Ramp induced currents simultaneously measured with membrane capacitance, also scaled to linear capacitance. Insert: top traces show large nonlinear displacement currents extracted with P/-5 protocol, subtraction holding potential set to +50 mV. Voltage protocol (holding potential 0 mV) shown below traces.

#### Inferred lipid packing within and around prestin

While detergent micelles have distinct properties from the lipid bilayer, we note an intimate relationship between the protein and micelle (**Figure 5**). Compared to other members of the extended transporter family, the two protomers of the prestin dimer show minimal interactions between the transmembrane domains as is the case with Slc26a9. Thus, the space between the membrane domains of the two protomers is filled by the detergent micelle (**Figure 5E**). Moreover, with the caveat that micelles may not be analogous to a bilayer, we note the distance between the inner and outer “leaflets” of the micelle tend to vary across the protein’s landscape (**Figure 5 A**,**B**,**C**,**D**). **Figure 5, Supplement 1** shows a corresponding variation in prestin’s surface hydrophobicity across its transmembrane domain. These data raise a number of issues pertaining to lipid effects on the protein’s function. For example, perhaps the variations in “leaflet” thickness is a reason for the shallow voltage dependence of prestin’s NLC. We consider the influence of the surrounding lipid bilayer below.

**Figure 5.**
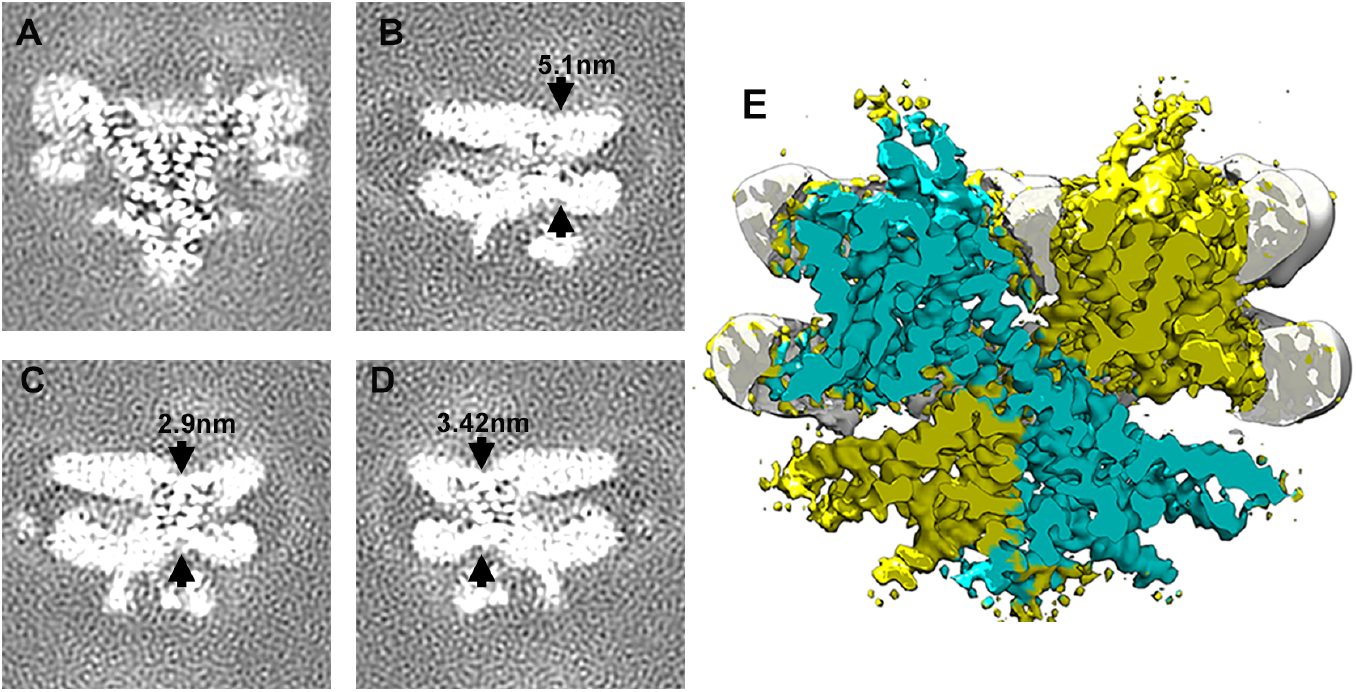
(**A, B**,**C**,**D**) Gray scale slices through the 3D reconstruction of prestin showing variations in the thickness of the micelle belt surrounding the protein. The distances measured within the density of the micelle belt are indicated by arrows in each panel. (**E**) Slice view of the cryo-EM reconstruction of prestin showing the micelle belt (in white) surrounding the transmembrane domains of prestin (in cyan and yellow).

## Discussion

We found that the overall molecular architecture of prestin is quite similar to the previously reported cryo-EM structures of full-length human SLC26A9 (Walter et al., 2019; Chi et al., 2020) and of a shorter version of mouse Slc26a9 protein (Walter et al., 2019; Chi et al., 2020), whose functions are to facilitate the transport of anions. Slc26a9 allows entry of the substrates (anions) from the cytosol through the so-called “inward-open” conformation, which shows an opening between the “core” and the “gate” domains as described in the solved cryo-EM structures (Walter et al., 2019; Chi et al., 2020). Intriguingly, we found that the membrane-spanning portion of prestin does not conform to the “inward-open” conformation of Slc26a9. Rather, prestin has a more compact conformation, closer to the “inward-open/intermediate” conformation of Slc26a9 (PDB ID, 6RTF), which shows the cytosol-exposed space between the “core” and the “gate” domains to be tighter. We reason that our prestin conformation solved by cryo-EM represents the contracted state of the protein at 0 mV.

We have compared our prestin cryo-EM structure to the predicted structure by the AlphaFold algorithm and find that despite overall similar topology, the pair-wise C_alpha_ RMSD calculated over the number of residues resolved in the cryo-EM structure is 7.3 Å.

### Key functional implications of prestin structure

The essential work of the voltage-dependent protein prestin dwells at frequencies where mammals can hear, driven by OHC AC receptor potentials. That is, the protein works in the kilohertz range of conformational change, with high-frequency measures of NLC reporting indirectly on those motions (Santos-Sacchi and Tan, 2020). Cryo-EM structures of prestin, which necessarily define one or more steady state conformations of the protein, may tell us little about its high frequency physiology. Indeed, we cannot be sure that any identified state is actually occupied during high frequency voltage stimulation, where molecular interactions, e.g. at the lipid-protein interface (see **Figure 5**), may be influenced by rate (frequency) itself. The stretched-exponential nature of prestin’s NLC likely reflects such interactions (Santos-Sacchi et al., 2009). Nevertheless, static structures can inform on some key questions concerning prestin’s molecular behavior.

Given the anion binding pocket structural features (not assumed from other family members), we can unequivocally assign relevance to prior and present mutational perturbations. Thus, we present data here that Cl^-^ does not function as prestin’s extrinsic voltage sensor, evidenced by the loss of anion sensitivity with structurally driven mutations in the anion binding site. Instead, prestin possesses charged residues important for voltage sensing, akin to other voltage sensing membrane proteins (Bezanilla, 2008; Bai et al., 2009). The distribution of twelve charged residues that sense voltage in prestin has important implications. Seven of the 12 residues are located in the gate/dimerization domain in TMs 5,6 and 12 that are modeled to move minimally in the elevator model (Bai et al., 2009; Drew and Boudker, 2016; Ficici et al., 2017). Notably, all but two of the 12 residues lie in the intracellular halves of the TM domains or in the intracellular loop connecting these TM domains (**Supplemental Figure 6**). In contrast, all but one of the five charged residues that have no effect on voltage sensing lie close to the extracellular halves of the TM domain or in the loops connecting these TM domains. This ineffectual group includes a charged residue in TM 3 (R150) that is modeled to move significantly in the transporter cycle (Bai et al., 2009; Drew and Boudker, 2016; Ficici et al., 2017). Together, these data suggest that contrary to functional expectations based solely on structural similarity between prestin and Slc26a9, electromechanical behavior in prestin is fundamentally different to transporter movements and is concentrated in proximity to the intracellular opening.

The wide dispersion of residue charges in several transmembrane domains that contribute to NLC and the inferred uneven lipid packing may underlie prestin’s shallow voltage dependence. Indeed, the influence of lipids on prestin performance is well documented (Santos-Sacchi and Wu, 2004; Sfondouris et al., 2008; Fang et al., 2010; Zhai et al., 2020). In agreement with the area motor model of prestin activity (Santos-Sacchi, 1993; Iwasa, 1994), we previously identified an augmentation of linear capacitance (ΔC_sa_) during hyperpolarization in prestin-transfected cells, our inducible prestin cell line (**Figure 3A**) and OHCs that likely reflects an increase in membrane surface area and membrane thinning accompanying movement of prestin into the expanded state (Santos-Sacchi and Navarrete, 2002; Santos-Sacchi and Song, 2014). Thus, we conclude that the compact state that we observe structurally corresponds to minimal membrane surface area and maximal membrane thickness. The converse would be expected for the expanded state. In this regard, although salicylate blocks NLC and eM, the effect it has on changes in linear capacitance (ΔC_sa_) indicates that it does not produce a natural state of prestin that is normally driven by voltage. That is, in the presence of salicylate the area occupied by prestin, as indicated by ΔC_sa_, is doubled that produced in its absence when driven by voltage (Santos-Sacchi and Navarrete, 2002; Santos-Sacchi and Song, 2014). Thus, we might expect that direct voltage-driven conformational changes in prestin could differ from anion-induced steady state cryo-EM structures, as we have exploited with high Cl^-^ levels here. Interestingly, we have recently shown that the ΔC_sa_ component of membrane capacitance exists only in the real (capacitive) part of complex NLC, not in the imaginary (conductive) part, ostensibly linking it directly to membrane bilayer influence rather than prestin charge movement (Santos-Sacchi et al., 2021).

We have determined that prestin is dimeric as our previous studies have suggested (Navaratnam et al., 2005; Bian et al., 2013), although earlier reports asserted that prestin functions as a tetramer (Wang et al., 2010; Hallworth and Nichols, 2012). In Slc26a9, three features were identified as important for dimerization and likely are pertinent for prestin considering the proteins remarkable similarity. These include **1**) membrane interactions between individual protomers exemplified by the valine zipper in TM14, **2**) interactions between the C terminal STAS domain of one protomer and the TM domain of the other and **3**) an antiparallel beta-strand between the N terminus of both protomers. Prestin shows each of these same interactions with the valine zipper being replaced by leucine and isoleucine residues. The anti-parallel beta-sheet between the N terminal residues 15-20 of each protomer in prestin is likely critical for dimerization. Indeed, sequential deletion of residues 11-21 resulted in progressive loss of NLC, and loss of FRET signal confirming the loss of dimerization (Navaratnam et al., 2005). These data also argue that dimerization is critical for NLC, just as dimerization is important for transporter function in UraA (Lu et al., 2011; Yu et al., 2017).

The C-terminus of prestin has high homology to the previously determined X-ray crystal structure of the C-terminal STAS domain (PDB ID, 3LLO) lacking the unstructured loop (Pasqualetto et al., 2010). Thus, the many interpretations made by the Battistutta group are likely to therefore apply, including confirmation in our structure of the orientation of the STAS domain to the transmembrane domain, and the importance of the alpha 5 helix in stabilizing the core beta sheets suggested by truncation experiments (Navaratnam et al., 2005; Zheng et al., 2005). A significant difference was in the first alpha helix that is parallel to the second alpha helix as in bacterial ASA (anti-sigma factor antagonistic) proteins (Aravind and Koonin, 2000), and deviated by a 30° angle in the crystal structure (Pasqualetto et al., 2010). This is likely due to the unstructured loop at the end of the first alpha helix that was lacking in the crystal structure.

Finally, structural details of the C-terminal STAS domain can shed light on modulation of prestin by binding partners, for example, calcium/calmodulin (Lolli et al., 2015). Recently, calmodulin has been shown to bind to the STAS domain (Costanzi et al., 2021). Such interactions may have physiological significance, since it was reported that Ca^2+^/calmodulin shifts the operating voltage of prestin, namely V_h_ (Keller et al., 2014). However, it was subsequently shown that shifts in V_h_ were not due to direct action on prestin, but rather resulting from indirect tension effects on prestin due to OHC swelling (Song and Santos-Sacchi, 2015). The influence of the STAS domain on prestin’s voltage-dependent activity remains an open question.

### Summary

OHC electromotility was discovered in 1985 (Brownell et al., 1985; Kachar et al., 1986), and 15 years later prestin was identified as the protein responsible for the OHC’s unique role in cochlear amplification (Zheng et al., 2000; Ludwig et al., 2001; Santos-Sacchi et al., 2001). During the intervening years enormous detail into the protein’s function has been obtained, see (Santos-Sacchi et al., 2017). Recent homology modelling of prestin (Gorbunov et al., 2014), based on presumed similarity to other family members, and confirmed in our structural data, has moved us closer to understanding prestin’s electromechanical behavior. Indeed, the cryo-EM solution we provide here will permit us to rigorously interpret past studies and design new experiments to more fully understand prestin’s role in mammalian hearing. Given the remarkable similarity of prestin structure revealed in this present study to that of the inside open/ intermediate state of Slc26a9, key questions remain as to why the two proteins differ in their function. Indeed, established functional observations on prestin may have already identified key differences, for example, the observed negative cooperativity among the densely packed proteins interacting through membrane lipids (10,000/μm^2^ in OHCs) (Santos-Sacchi et al., 1998; Zhai et al., 2020), and subplasmalemmal cytoskeletal interactions with prestin (Bai et al., 2010). Though key, prestin likely is a partner in the machinery that boosts our hearing abilities. Imperative in the overall effort to understand prestin is the need for alternative structures of prestin and other family members that are evoked by appropriate physiological stimuli, e.g., voltage in the case of prestin.

## Materials and Methods

### Prestin Expression and Purification

The full-length prestin from gerbil (*Meriones unguiculatus, Genbank accession number* AF230376) was purified from a tetracycline inducible stable HEK 293 cell line (Bian et al., 2010). In establishing this cell line (16C), full length gerbil prestin cDNA (a gift from J. Zheng and P. Dallos) tagged at its C-terminus with enhanced yellow fluorescent protein (EYFP) was inserted into the multiple cloning site of pcDNA4/TO/*myc*-HisC that allowed purification using Ni affinity.

Cells were grown in DMEM media supplemented with 1 mM L-glutamine, 100 U ml^−1^ penicillin/streptomycin, 10% FBS and 1 mM sodium pyruvate. 4 μg/ml of blasticidin and 130 μg/ml of zeocin were supplemented in the growth media to maintain prestin expression. Cells were harvested 48 hours after tetracycline (1 μg/ml) was added to the cell growth medium to induce prestin expression.

Cell pellets from 20 T175 Flasks were harvested by centrifugation at 1,000g for 10 minutes, washed with PBS, then resuspended in 5ml of resuspension buffer (25 mM HEPES, pH 7.4, 200 mM NaCl, 5% glycerol, 2mM CaCl_2_, 10 µg/ml^-1^ DNase I and 1 protease inhibitors (cOmplete EDTA-free, Roche) for each gram of pellet. 2% (wt/vol final concentration) digitonin (Anatrace) powder was directly dissolved in the cell resuspension, and the mixture was incubated for 1.5 hr under gentle agitation (rocking) at 4°C. Insoluble material was removed by centrifugation at 160,000 g for 50 minutes (Beckman L90-XP ultracentrifuge with a 50.2 Ti rotor). The supernatant was passed through a 0.45 µm filter. 10 mM imidazole (final concentration) and 1 ml Ni-NTA resin (Qiagen) pre-washed in 25mM Hepes, pH 7.4, 200 mM NaCl, 5% glycerol, 2mM CaCl_2_, 0.02% GDN (synthetic digitonin substitute glyco-diosgenin, Antrace) were added to the filtered supernatant and incubated with end over end rocking agitation at 4 °C for 2 hours. The resin was collected using a bench Eppendorf microcentrifuge and washed sequentially with high salt buffer Buffer A (25 mM HEPES, pH 7.4, 500 mM NaCl, 5% glycerol, and 0.02% GDN) and 25ml Buffer B (25 mM HEPES, pH 7.4, 200 mM NaCl, 10 mM imidazole, 5% glycerol, and 0.02% GDN). The protein was eluted in 1.5ml Buffer C (25 mM HEPES, pH 7.4, 200 mM NaCl, 250 mM imidazole, 5% glycerol, and 0.02% GDN). The 1.5 ml eluted protein was concentrated to 500 µl, passed through a 0.22 µm filter and loaded onto a FSEC column (Superdex 200 Increase 10/300 GL column, on a Shimadzu FPLC system) equilibrated with gel-filtration buffer (10 mM HEPES, 200 mM NaCl, 0.02% GDN, pH 7.4). Two 0.5 ml fractions corresponding to the fluorescent (excitation 488 nm, emission 535 nm) peak and A280 peak were collected and concentrated using an Amicon Ultra centrifugal filter with a molecular weight cutoff 100 KDa and used for freezing grids.

### Sample Preparation and Data Acquisition

An aliquot of four microliters of purified prestin (at a concentration of approximately 2 mg/ml) was applied to glow-discharged Quantifoil holey carbon grids (Gold R2/1; 200 mesh) overlaid with an additional 2-nm carbon layer (Electron Microscopy Sciences). The grids were blotted for 3-5 s and plunge-frozen in liquid ethane using a Vitrobot Mark IV (FEI) instrument with the chamber maintained at 10°C and 100% humidity.

Cryo-EM micrograph movies were recorded using the SerialEM software on a Titan Krios G2 transmission electron microscope (Thermo Fischer/FEI) operated at a voltage of 300 kV and equipped with a K3 Summit direct electron detector (Gatan, Pleasanton, CA). A quantum energy filter with a 20 eV slit width (Gatan) was used to remove the inelastically scattered electrons. In total 4,680 dose-fractionated super-resolution movies with 36 images per stack were recorded. The cryo-EM movies were recorded with a defocus varied from –1.15 to –2.15 µm at a nominal magnification of 81,000x (corresponding to 0.534 Å per physical pixel). The counting rate was 17.5 e^-^/physical pix/s. The total exposure time was 3.6 s per exposure with a total dose of ∼ 54 e^-^/Å^2^.

### Data Processing

Data processing was carried out with Relion 3.1 (Zivanov et al., 2018) except as noted. Movies frames were gain-normalized and motion-corrected using MotionCor2 with a binning factor of 2 and dividing micrographs into 4×4 patches. The dose-weighted, motion corrected micrographs (the sum of 36 movie frames) were used for all image processing steps except for defocus determination. The contrast transfer function (CTF) calculation was performed with CTFFIND4.1 (as implemented in Relion3.1) on movie sums which were motion-corrected but not dose-weighted. About 3,000 particles were manually picked and subjected to 2D reference-free classification in Relion 3.1 (Zivanov et al., 2018).

Classes showing good signal (representing ∼1,100 particles) were chosen as references for automated particle picking in Relion 3.1, yielding a data set of ∼1,377,109 particles. Several rounds of 2D and 3D classification (carried out without application of symmetry) were used to remove unsuitable particles, leaving 122,754 particles that were used for structural determination with imposed C2 symmetry in Relion 3.1. Bayesian polishing (Zivanov et al., 2018), followed by per-particle CTF refinement, 3D auto-refinement, micelle density subtraction and post-processing generated a map which had an estimated resolution of ∼ 4 Å according to the Fourier shell correlation (FSC) =0.143 criterion.

### Model Building

The SLC26a9 cryo-EM intermediate structure (PDB ID, 6RTF) which is a poly-alanine trace) was rigid-body docked into the prestin cryo-EM map and fitted using Chimera (Pettersen et al., 2004). Next, backbone was real space-refined in Phenix (Adams et al., 2010) and adjusted in COOT (Emsley et al., 2010) by manually going through the entire protomer chains. Sequence assignment was guided mainly by bulky residues such as Phe, Tyr, Trp, and Arg, and secondary structure predictions. Side chains in areas of the map with insufficient density were left as alanine. The model was refined through several rounds of model building in COOT and real-space refinement in PHENIX (Adams et al., 2010) with secondary structure and geometry restraints. Details on the statistics of cryo-EM data collection and structure determination are presented in Supplementary Table 1. Figures were prepared using Chimera (Pettersen et al., 2004) and ChimeraX (Pettersen et al., 2021).

### Electrophysiological recording

Recordings were made of transiently transfected CHO (or HEK cells 48 hours after tetracycline induction) using a whole-cell configuration at room temperature using an Axon 200B amplifier (Molecular Devices, Sunnyvale, CA), as described previously. Cells were recorded 48– 72 h after tetracycline induction or transfection (Fugene (Promega) according to the manufacturer’s instructions) to allow for stable measurement of current and NLC. Mutations were introduced in gerbil prestin YFP using the QuickChange Mutagenesis Kit (Agilent) according to the manufacturer’s instructions. The standard bath solution components were (in mM): 100 NaCl (TEA)-Cl 20, CsCl 20, CoCl_2_ 2, MgCl_2_ 2, Hepes 5, pH 7.2. In addition, 20mM NaSCN was substituted for current recordings with cells transiently transfected with Slc26a9. The pipette solution contained (in mM): NaCl 100, CsCl 20, EGTA 5, MgCl_2_ 2, Hepes 10, pH 7.2. Osmolarity was adjusted to 300 ± 2 mOsm with dextrose. After whole cell configuration was achieved in extracellular NaCl a ramp protocol recorded to confirm baseline NLC and currents. Pipettes had resistances of 3-5 MΩ. Gigohm seals were made and stray capacitance was balanced out with amplifier circuitry prior to establishing whole-cell conditions. A Nikon Eclipse E600-FN microscope with 40× water immersion lens was used to observe cells during voltage clamp. Data were low pass filtered at 10 kHz and digitized at 100 kHz with a Digidata 1320A.

Command delivery and data collections were carried out with a Windows-based whole-cell voltage clamp program, jClamp (Scisoft, East Haven, CT), using a Digidata 1322A (Axon Instruments). A continuous high-resolution 2-sine voltage command was used, cell capacitance and current being extracted synchronously. In order to extract Boltzmann parameters, capacitance-voltage data were fit to the first derivative of a two-state Boltzmann function.

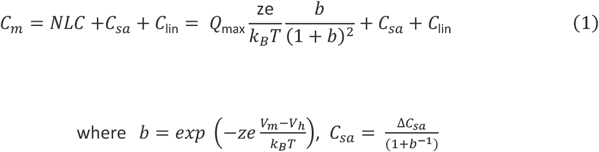

Q_max_ is the maximum nonlinear charge moved, V_h_ is voltage at peak capacitance or equivalently, at half-maximum charge transfer, V_m_ is membrane potential, *z* is valence, C_lin_ is linear membrane capacitance, e is electron charge, *k*_*B*_ is Boltzmann’s constant, and T is absolute temperature. C_sa_ is a component of capacitance that characterizes sigmoidal changes in specific membrane capacitance, with ΔC_sa_ referring to the maximal change at very negative voltages (Santos-Sacchi and Navarrete, 2002; Santos-Sacchi and Song, 2014). For fits of NLC in transiently transfected cells, C_sa_ was not included. *Q*_sp_ the specific charge density, is the total charge moved (*Q*_max_) normalized to linear capacitance. Voltages were corrected for series resistance voltage drop. Separately, in specific experiments gating currents were also determined using voltage steps (50 ms duration) from −100 mV to 150 mV, with 20 mV step increments. Where anions were substituted, local perfusion of the cells were estimated to give rise to small junctional potentials (JPCalc function in pClamp). Since these numbers were small no corrections were made to the IV plots. For solution perfusion experiments we used a QMM perfusion system (ALA Scientific, Instruments, Westbury, NY). The manifold’s output tip was 200 μm placed 1 mm from the cell, and the flow rate increased by an applied pressure of approximately 20 kPa. Statistical analysis was done with SAS software (SAS Institute Inc, NC).

## Acknowledgements

This research was supported by NIH-NIDCD R01 DC016318 (JSS) and R01 DC008130 (JSS, DN).

## Legends

**Figure 1 Supplement 1.**
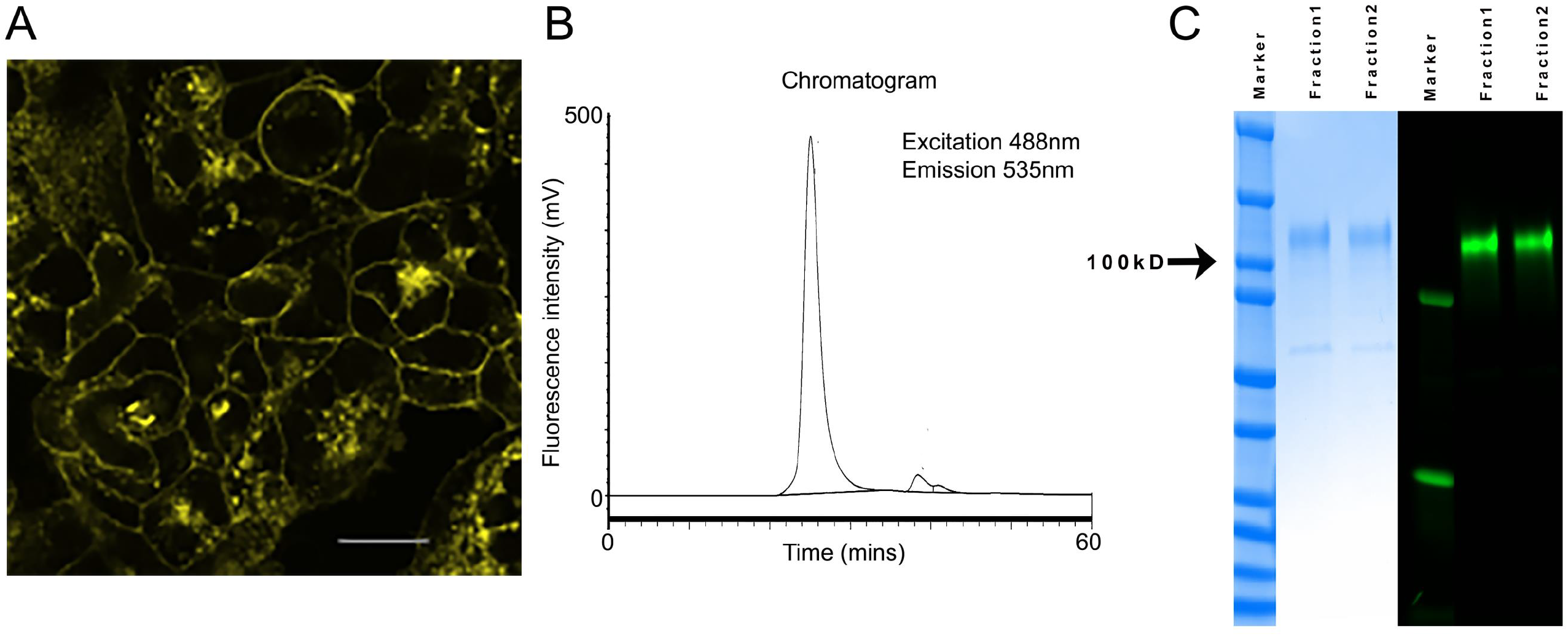
(**A**). The figure shows a fluorescence image of HEK 293 cells 48 hours after tetracycline induction with prestin YFP expressed in the plasma membrane of cells. Scale bar is 10 microns. (**B**) Fluorescence Size Exclusion Chromatogram (FSEC) of purified prestin YFP injected into a Superdex 200 Increase 10/300 GL Column. Two fractions corresponding to the single monodisperse peak were collected and used for cryo-EM analysis. (**C**). Coomasie blue staining (left three lanes) and fluorescence in gel imaging of the two fractions obtained from FSEC confirmed the presence of purified prestin-YFP (∼110 kDa). The position of the 100 kDa molecular weight marker is indicated.

**Figure 1 Supplement 2.**
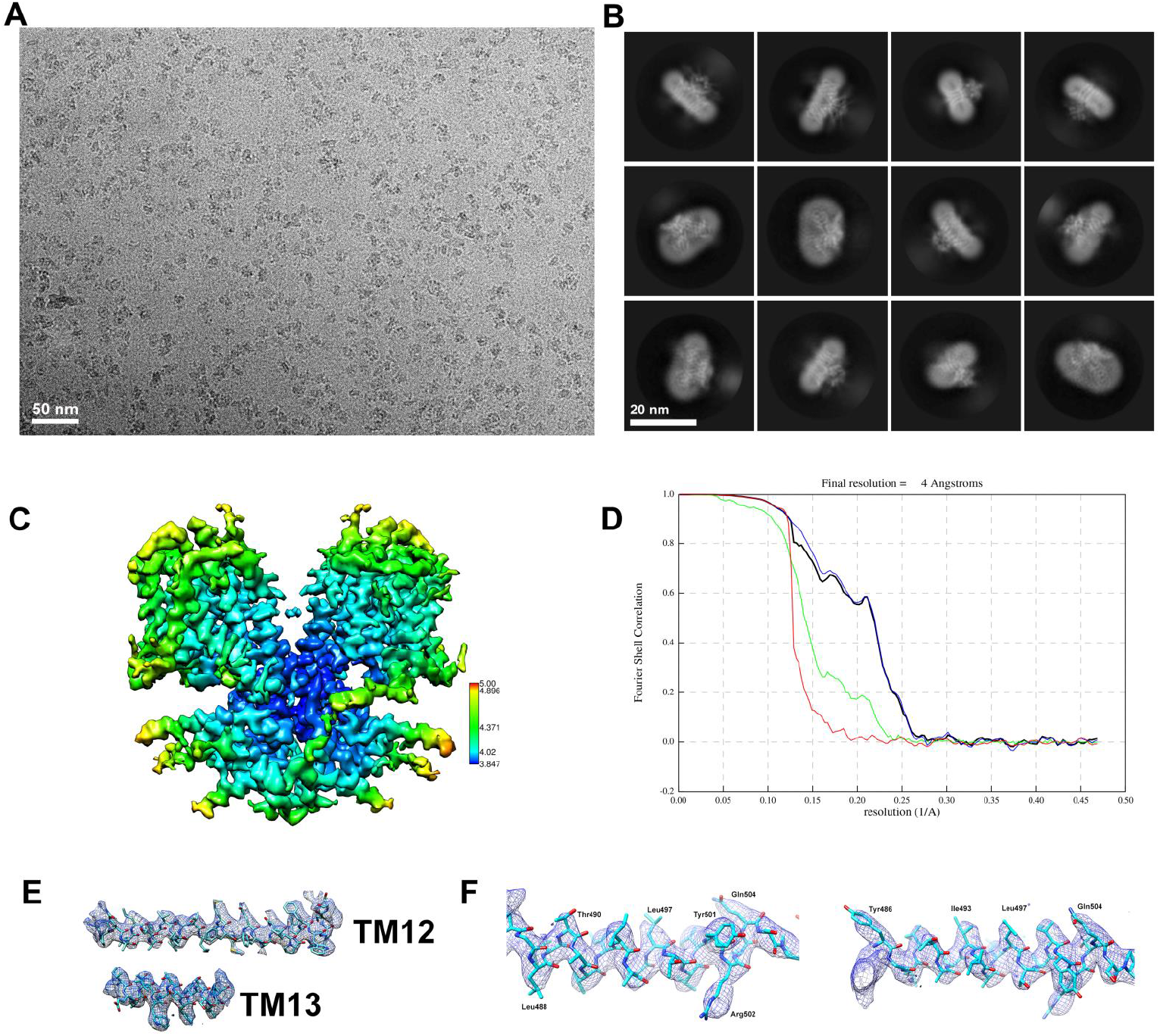
Cryo-EM imaging of prestin. (**A**) representative cryo-EM micrograph; (**B**) Selected reference-free 2D class averages; (**C**) Cryo-EM density map of prestin colored by local resolution. (D) Gold standard FSC curves from RELION indicate that the cryo-EM structure of prestin has a nominal resolution of 4.0 Å at FSC=0.143. (**E**) Examples of the fit of the prestin model into the density map. To the left, density and model for the TM12 and TM13 segments. To the right, representative density with the model for the L488-Q504 residues segment.

**Figure 2, Figure Supplement 1.**
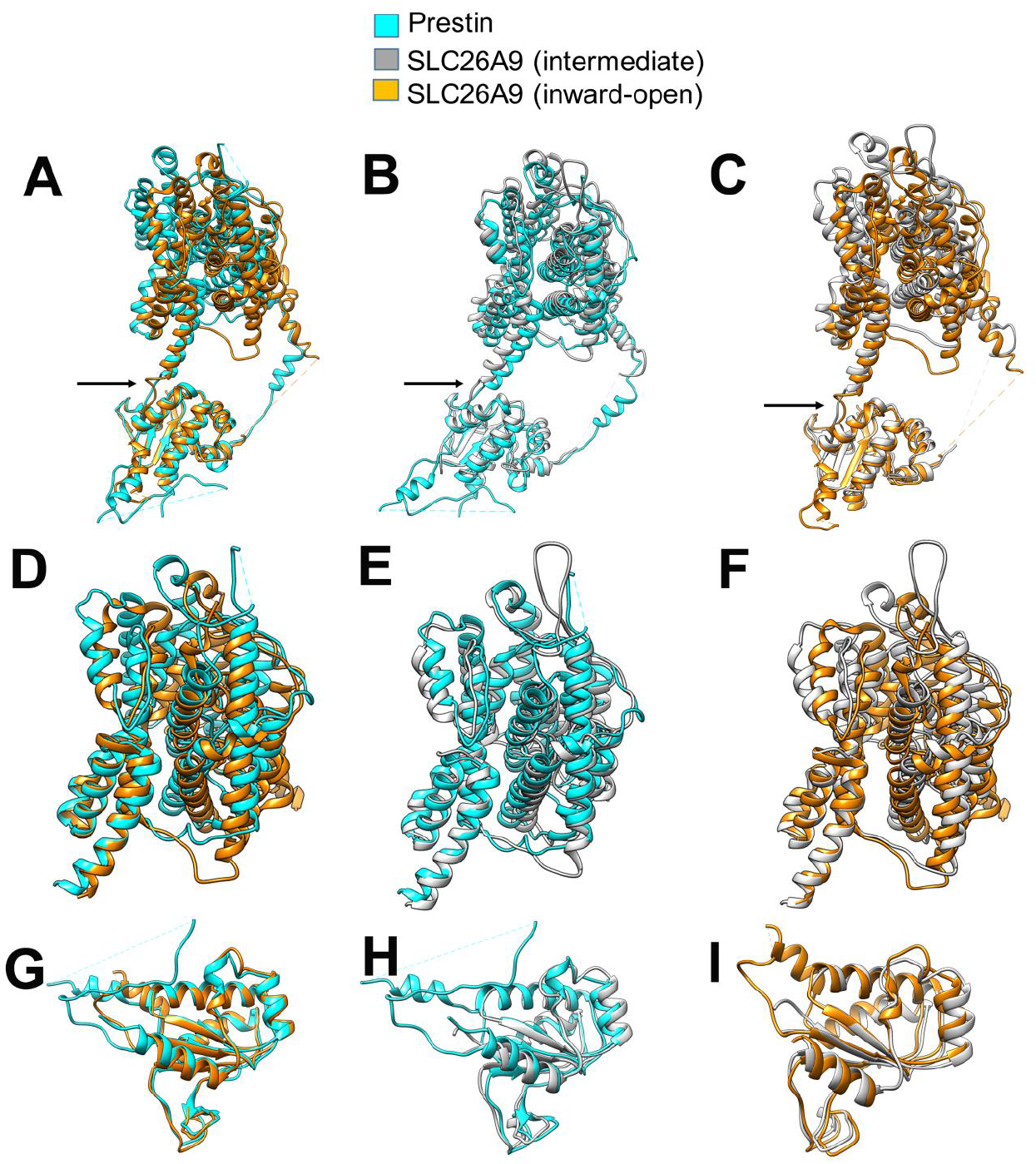
(**A**,**B**,**C**) Superposition of the prestin monomer structure with the Slc26a9 monomer structure. The monomers have been superposed through a segment within the STAS domain (Gln504-Ala578 within the prestin’s STAS domain). Arrow indicates a possible hinge at the connection between the cytosolic and membrane domain. (**D**,**E**,**F**) Superposition of the separated transmembrane domain of prestin with the separated transmembrane domains of Slc26a9 in “inward-open” (PDB ID, 7CH1, in orange) and “intermediate” conformations(PDB ID, 6RTF, in grey). (**G**,**H**,**I**) superposition of the separated STAS domain of prestin with the separated STAS domains of Slc26a9 in “inward-open” and “intermediate” conformations.

**Figure 2, Figure Supplement 2.**
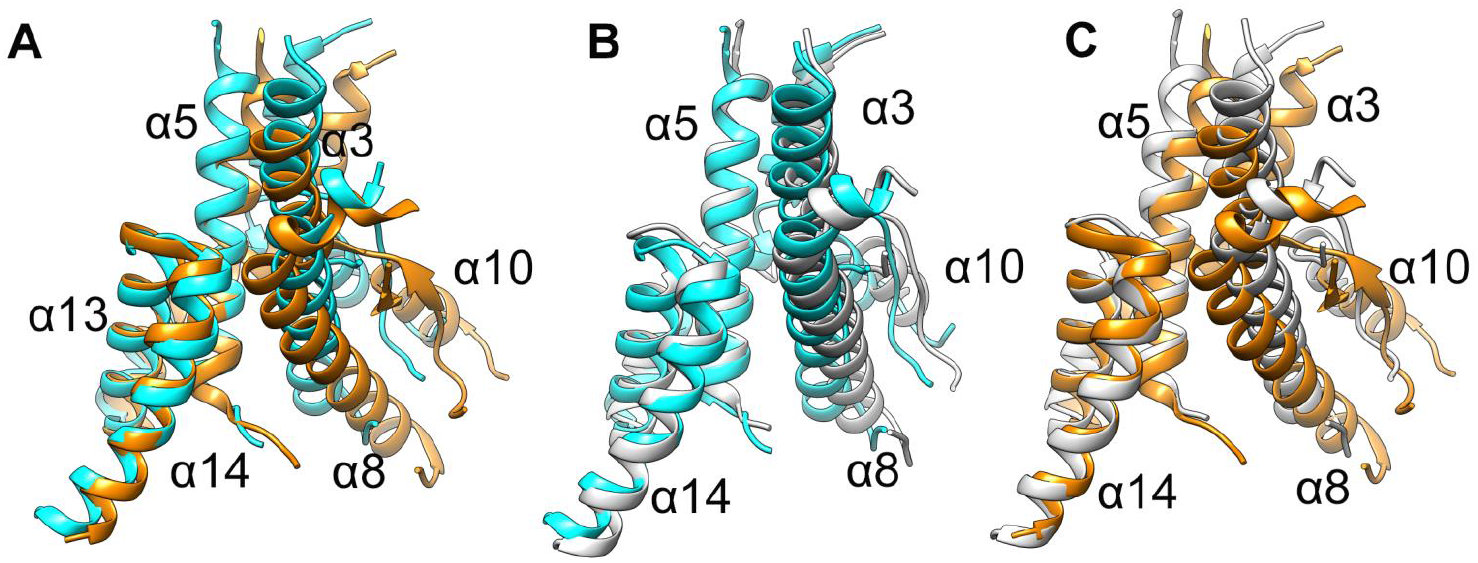
(**A**,**B**,**C**) Close-up view of the TM13, TM14 and TM8 helices from the superposition of the transmembrane domain of prestin with the transmembrane domains of Slc26a9 (PDB ID, 7CH1, in orange ribbon and PDB, ID, 6RTF, in grey ribbon). TM8 has different orientations with respect to TM13 and TM14 in prestin and Slc26a9.

**Figure 4 Supplement 1.**
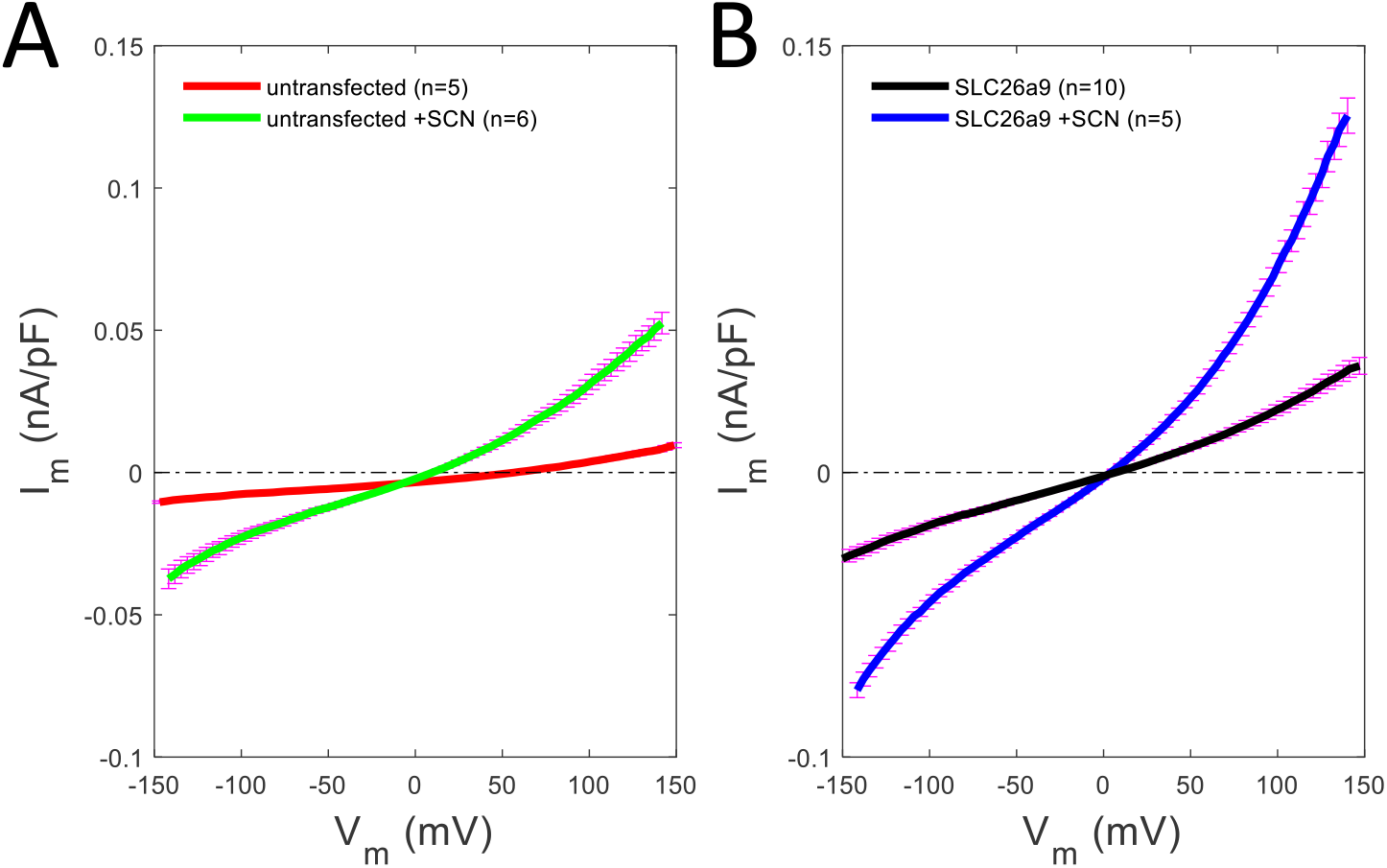
(**A**) Effect of SCN^-^ on the magnitude of ramp induced current in un-transfected CHO cells. A slight increase in current is observed with SCN^-^. (**B**) In Slc26a9 transfected cells, larger currents are observed without and with SCN^-^, indicating successful delivery of the construct to the membrane.

**Figure 5 Supplement 1.**
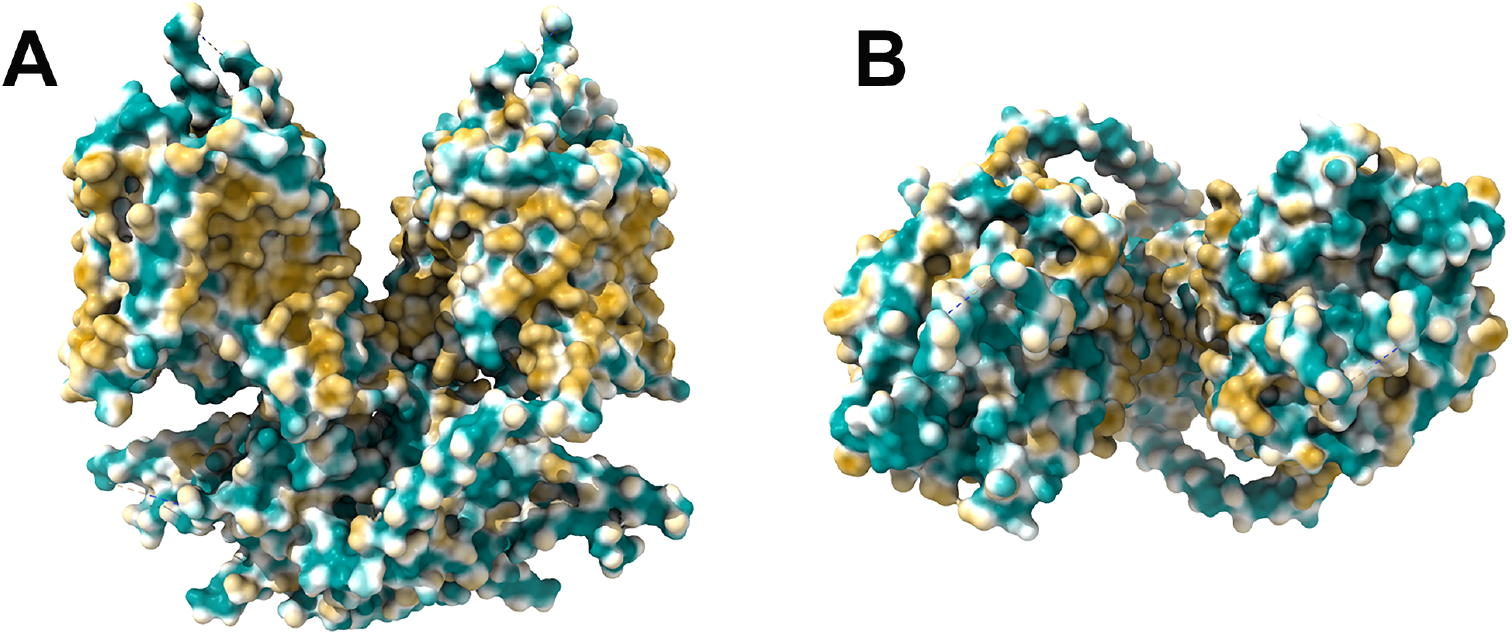
(**A**,**B**) Representation of hydrophobicity on the prestin surface with the most extensive hydrophobic patches located within the TM domains of the protein (in brown). Hydrophilic residues are shown in green.

**Supplemental Figure 6.**
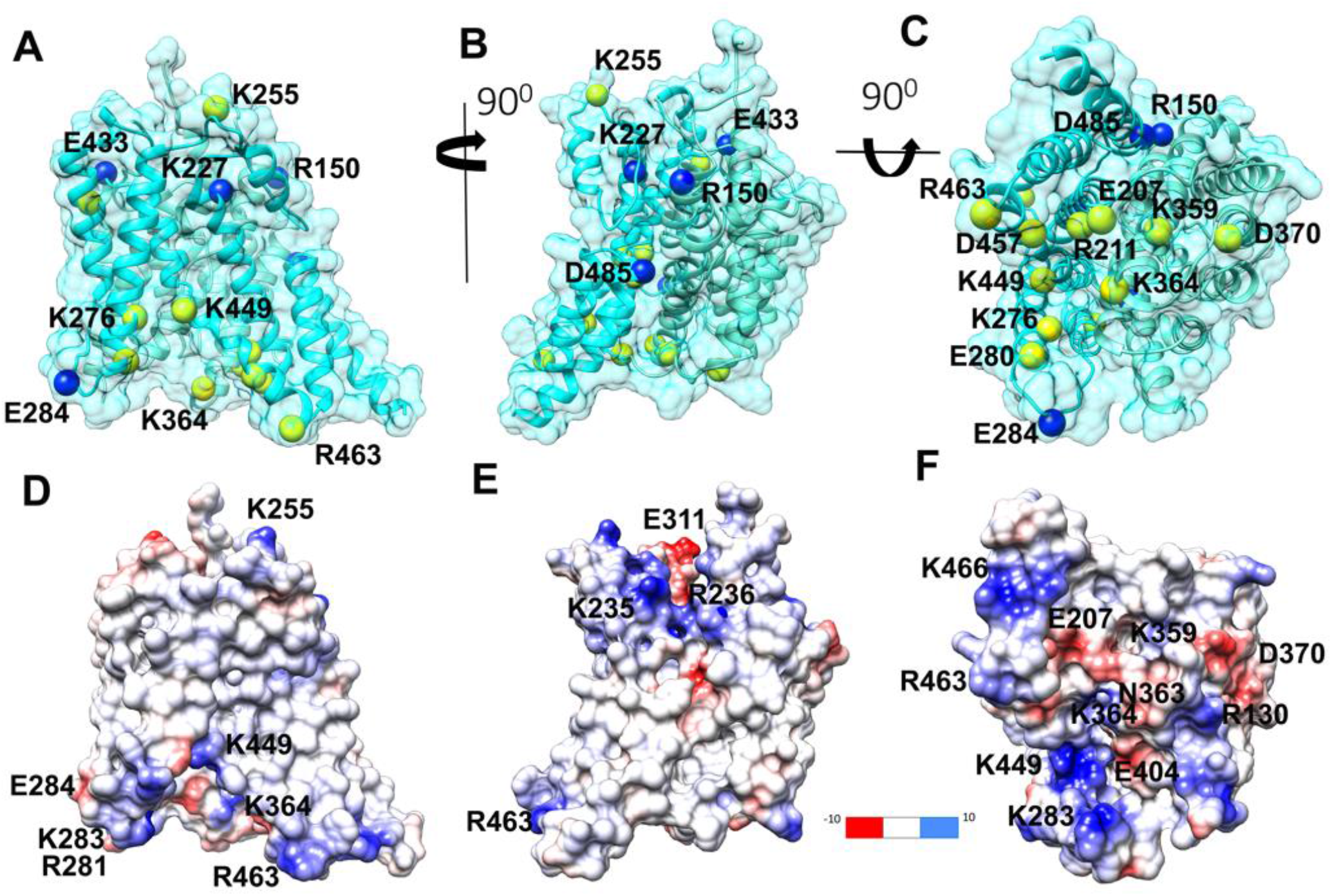
Mapping of charged residues important for the nonlinear capacitance onto the prestin surface. (**A, B, C**) Three views of the transmembrane domain of prestin monomer shown in transparent surface representation. The position of charged residues (R130, E207, R211, K255, K276, E280, K359, K364, D370, K449, D457, R463) which are important for the NLC is highlighted by yellow spheres. Mapping of these functionally important residues onto the prestin surface shows that they cluster preferentially within the cytosolic exposed surface. Positions of charged residues (R150, K227, E284, E433, D485) which are not important for the NLC are highlighted by blue spheres. (**D**,**E**,**F**) Electrostatic surface of the prestin’s TM domain shown in the same orientations as the views of the prestin’s TM surface in panels (**A**,**B**,**C**). Charged residues are indicated (red negative, blue positive).

**Table S1.**
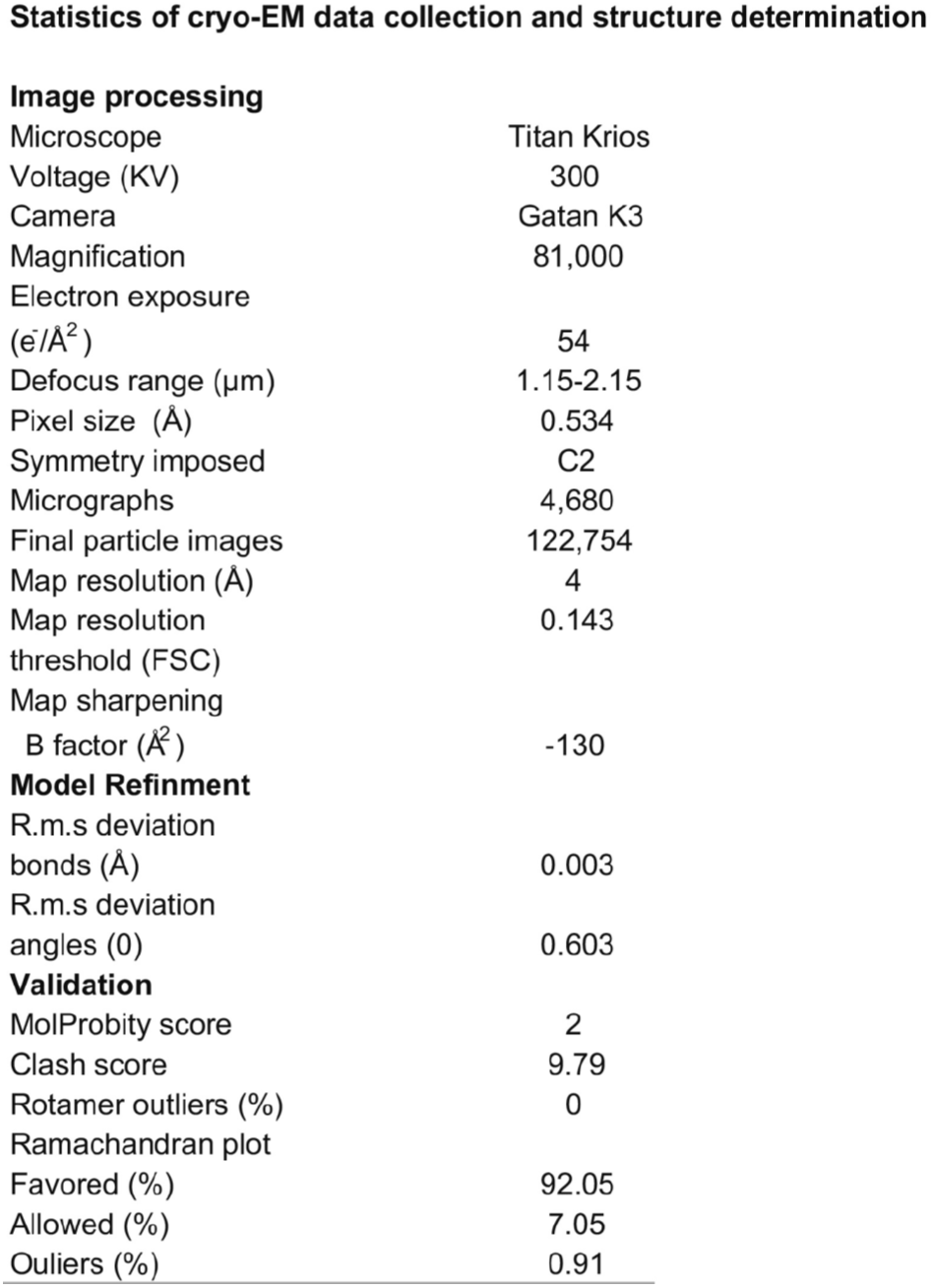
Cryo-EM data collection, refinement and validation statistics

|  |  |
| --- | --- |
| Microscope | Titan Krios |
| Voltage (KV) | 300 |
| Camera | Gatan K3 |
| Magnification | 81,000 |
| Electron exposure<br>($\text{e}^-/\text{\AA}^2$) | 54 |
| Defocus range ($\mu\text{m}$) | 1.15-2.15 |
| Pixel size ($\text{\AA}$) | 0.534 |
| Symmetry imposed | C2 |
| Micrographs | 4,680 |
| Final particle images | 122,754 |
| Map resolution ($\text{\AA}$) | 4 |
| Map resolution<br>threshold (FSC) | 0.143 |
| Map sharpening<br>B factor ($\text{\AA}^2$) | -130 |
| **Model Refinement** |  |
| R.m.s deviation<br>bonds ($\text{\AA}$) | 0.003 |
| R.m.s deviation<br>angles ($^\circ$) | 0.603 |
| **Validation** |  |
| MolProbity score | 2 |
| Clash score | 9.79 |
| Rotamer outliers (%) | 0 |
| Ramachandran plot<br>Favored (%) | 92.05 |
| Allowed (%) | 7.05 |
| Outliers (%) | 0.91 |

## Notes

### Competing Interest Statement

The authors have declared no competing interest.

### Summary of Updates

Significant revision to introduction, results and discussion

## References

Adams PD, Afonine PV, Bunkoczi G, Chen VB, Davis IW, Echols N, Headd JJ, Hung LW, Kapral GJ, Grosse-Kunstleve RW, McCoy AJ, Moriarty NW, Oeffner R, Read RJ, Richardson DC, Richardson JS, Terwilliger TC, Zwart PH (2010) PHENIX: a comprehensive Python-based system for macromolecular structure solution. Acta Crystallogr D Biol Crystallogr 66:213–221.

Aravind L, Koonin EV (2000) The STAS domain - a link between anion transporters and antisigmafactor antagonists. Curr Biol 10:R53–55.

Ashmore J (2005) Prestin is an electrogenic anion transporter. 42ndWorkshop on Inner Ear Biology Sept.:138.

Ashmore JF (1990) Forward and reverse transduction in the mammalian cochlea. Neurosci Res Suppl 12:S39–S50.

Bai JP, Surguchev A, Montoya S, Aronson PS, Santos-Sacchi J, Navaratnam D (2009) Prestin’s anion transport and voltage-sensing capabilities are independent. Biophys J 96:3179–3186.

Bai JP, Surguchev A, Ogando Y, Song L, Bian S, Santos-Sacchi J, Navaratnam D (2010) Prestin surface expression and activity are augmented by interaction with MAP1S, a microtubule-associated protein. JBiolChem 285:20834–20843.

Bai JP, Moeini-Naghani I, Zhong S, Li FY, Bian S, Sigworth FJ, Santos-Sacchi J, Navaratnam D (2017) Current carried by the Slc26 family member prestin does not flow through the transporter pathway. Sci Rep 7:46619.

Bezanilla F (2008) How membrane proteins sense voltage. NatRevMolCell Biol 9:323–332.

Bian S, Navaratnam D, Santos-Sacchi J (2013) Real time measures of prestin charge and fluorescence during plasma membrane trafficking reveal sub-tetrameric activity. PLoS One 8:e66078.

Bian S, Koo BW, Kelleher S, Santos-Sacchi J, Navaratnam DS (2010) A highly expressing Tet-inducible cell line recapitulates in situ developmental changes in prestin’s Boltzmann characteristics and reveals early maturational events. AmJPhysiol Cell Physiol 299:C828–C835.

Brownell WE, Bader CR, Bertrand D, de Ribaupierre Y (1985) Evoked mechanical responses of isolated cochlear outer hair cells. Science 227:194–196.

Chang YN, Geertsma ER (2017) The novel class of seven transmembrane segment inverted repeat carriers. Biol Chem 398:165–174.

Chang YN, Jaumann EA, Reichel K, Hartmann J, Oliver D, Hummer G, Joseph B, Geertsma ER (2019) Structural basis for functional interactions in dimers of SLC26 transporters. Nat Commun 10:2032.

Chi X, Jin X, Chen Y, Lu X, Tu X, Li X, Zhang Y, Lei J, Huang J, Huang Z, Zhou Q, Pan X (2020) Structural insights into the gating mechanism of human SLC26A9 mediated by its C-terminal sequence. Cell Discov 6:55.

Costanzi E, Coletti A, Zambelli B, Macchiarulo A, Bellanda M, Battistutta R (2021) Calmodulin binds to the STAS domain of SLC26A5 prestin with a calcium-dependent, one-lobe, binding mode. J Struct Biol 213:107714.

Dallos P, Wu X, Cheatham MA, Gao J, Zheng J, Anderson CT, Jia S, Wang X, Cheng WH, Sengupta S, He DZ, Zuo J (2008) Prestin-based outer hair cell motility is necessary for mammalian cochlear amplification. Neuron 58:333–339.

Drew D, Boudker O (2016) Shared Molecular Mechanisms of Membrane Transporters. Annu Rev Biochem 85:543–572.

Emsley P, Lohkamp B, Scott WG, Cowtan K (2010) Features and development of Coot. Acta Crystallogr D Biol Crystallogr 66:486–501.

Fang J, Izumi C, Iwasa KH (2010) Sensitivity of prestin-based membrane motor to membrane thickness. Biophys J 98:2831–2838.

Ficici E, Faraldo-Gomez JD, Jennings ML, Forrest LR (2017) Asymmetry of inverted-topology repeats in the AE1 anion exchanger suggests an elevator-like mechanism. J Gen Physiol 149:1149–1164.

Gale JE, Ashmore JF (1994) Charge displacement induced by rapid stretch in the basolateral membrane of the guinea-pig outer hair cell. Proc R Soc Lond B Biol Sci 255:243–249.

Gorbunov D, Hartmann J, Renigunta V, Oliver D (2018) A glutamate scan identifies an electrostatic switch for prestin activity. Midwinter Meeting Abstracts of the Associatio for Research in Otolaryngology.

Gorbunov D, Sturlese M, Nies F, Kluge M, Bellanda M, Battistutta R, Oliver D (2014) Molecular architecture and the structural basis for anion interaction in prestin and SLC26 transporters. Nat Commun 5:3622.

Hallworth R, Nichols MG (2012) Prestin in HEK cells is an obligate tetramer. J Neurophysiol 107:5–11.

Iwasa KH (1993) Effect of stress on the membrane capacitance of the auditory outer hair cell. Biophys J 65:492–498.

Iwasa KH (1994) A membrane motor model for the fast motility of the outer hair cell. JAcoustSocAm 96:2216–2224.

Kachar B, Brownell WE, Altschuler R, Fex J (1986) Electrokinetic shape changes of cochlear outer hair cells. Nature 322:365–368.

Kakehata S, Santos-Sacchi J (1995) Membrane tension directly shifts voltage dependence of outer hair cell motility and associated gating charge. Biophys J 68:2190–2197.

Kakehata S, Santos-Sacchi J (1996) Effects of salicylate and lanthanides on outer hair cell motility and associated gating charge. J Neurosci 16:4881–4889.

Keller JP, Homma K, Duan C, Zheng J, Cheatham MA, Dallos P (2014) Functional regulation of the SLC26-family protein prestin by calcium/calmodulin. J Neurosci 34:1325–1332.

Kuwabara MF, Wasano K, Takahashi S, Bodner J, Komori T, Uemura S, Zheng J, Shima T, Homma K (2018) The extracellular loop of pendrin and prestin modulates their voltage-sensing property. J Biol Chem 293:9970–9980.

Lolli G, Pasqualetto E, Costanzi E, Bonetto G, Battistutta R (2015) The STAS domain of mammalian SLC26A5 prestin harbors an anion-binding site. Biochem J.

Lu F, Li S, Jiang Y, Jiang J, Fan H, Lu G, Deng D, Dang S, Zhang X, Wang J, Yan N (2011) Structure and mechanism of the uracil transporter UraA. Nature 472:243–246.

Ludwig J, Oliver D, Frank G, Klocker N, Gummer AW, Fakler B (2001) Reciprocal electromechanical properties of rat prestin: The motor molecule from rat outer hair cells. Proc Natl Acad Sci U S A 98:4178–4183.

Navaratnam D, Bai JP, Samaranayake H, Santos-Sacchi J (2005) N-terminal-mediated homomultimerization of prestin, the outer hair cell motor protein. Biophys J 89:3345–3352.

Oliver D, He DZ, Klocker N, Ludwig J, Schulte U, Waldegger S, Ruppersberg JP, Dallos P, Fakler B (2001) Intracellular anions as the voltage sensor of prestin, the outer hair cell motor protein. Science 292:2340–2343.

Pasqualetto E, Aiello R, Gesiot L, Bonetto G, Bellanda M, Battistutta R (2010) Structure of the cytosolic portion of the motor protein prestin and functional role of the STAS domain in SLC26/SulP anion transporters. J Mol Biol 400:448–462.

Pettersen EF, Goddard TD, Huang CC, Couch GS, Greenblatt DM, Meng EC, Ferrin TE (2004) UCSF Chimera--a visualization system for exploratory research and analysis. J Comput Chem 25:1605–1612.

Pettersen EF, Goddard TD, Huang CC, Meng EC, Couch GS, Croll TI, Morris JH, Ferrin TE (2021) UCSF ChimeraX: Structure visualization for researchers, educators, and developers. Protein Sci 30:70–82.

Rybalchenko V, Santos-Sacchi J (2003a) Allosteric modulation of the outer hair cell motor protein prestin by chloride. In: Biophysics of the Cochlea: From Molecules to Models (Gummer A, ed), pp 116–126. Singapore: World Scientific Publishing.

Rybalchenko V, Santos-Sacchi J (2003b) Cl-flux through a non-selective, stretch-sensitive conductance influences the outer hair cell motor of the guinea-pig. J Physiol 547:873–891.

Rybalchenko V, Santos-Sacchi J (2008) Anion control of voltage sensing by the motor protein prestin in outer hair cells. Biophys J 95:4439–4447.

Santos-Sacchi J (1991) Reversible inhibition of voltage-dependent outer hair cell motility and capacitance. J Neurosci 11:3096–3110.

Santos-Sacchi J (1993) Harmonics of outer hair cell motility. Biophys J 65:2217–2227.

Santos-Sacchi J, Navarrete E (2002) Voltage-dependent changes in specific membrane capacitance caused by prestin, the outer hair cell lateral membrane motor. Pflugers Arch 444:99–106.

Santos-Sacchi J, Wu M (2004) Protein-and lipid-reactive agents alter outer hair cell lateral membrane motor charge movement. JMembrBiol 200:83–92.

Santos-Sacchi J, Song L (2014) Chloride and Salicylate Influence Prestin-Dependent Specific Membrane Capacitance: Support for the Area Motor Model. J Biol Chem.

Santos-Sacchi J, Song L (2016) Chloride anions regulate kinetics but not voltage-sensor Qmax of the solute carrier SLC26a5. Biophys J 110:1–11.

Santos-Sacchi J, Tan W (2018) The Frequency Response of Outer Hair Cell Voltage-Dependent Motility Is Limited by Kinetics of Prestin. J Neurosci 38:5495–5506.

Santos-Sacchi J, Tan W (2019) Voltage Does Not Drive Prestin (SLC26a5) Electro-Mechanical Activity at High Frequencies Where Cochlear Amplification Is Best. iScience 22:392–399.

Santos-Sacchi J, Tan W (2020) Complex nonlinear capacitance in outer hair cell macro-patches: effects of membrane tension. Sci Rep 10:6222.

Santos-Sacchi J, Kakehata S, Takahashi S (1998) Effects of membrane potential on the voltage dependence of motility-related charge in outer hair cells of the guinea-pig. J Physiol 510 (Pt 1):225–235.

Santos-Sacchi J, Navarrete E, Song L (2009) Fast electromechanical amplification in the lateral membrane of the outer hair cell. Biophys J 96:739–747.

Santos-Sacchi J, Navaratnam D, Tan WJT (2021) State dependent effects on the frequency response of prestin’s real and imaginary components of nonlinear capacitance. Sci Rep 11:16149.

Santos-Sacchi J, Shen W, Zheng J, Dallos P (2001) Effects of membrane potential and tension on prestin, the outer hair cell lateral membrane motor protein. J Physiol 531:661–666.

Santos-Sacchi J, Song L, Zheng JF, Nuttall AL (2006) Control of mammalian cochlear amplification by chloride anions. J Neurosci 26:3992–3998.

Santos-Sacchi J, Navaratnam D, Raphael R, Oliver D (2017) The Cochlea Chapter 5: Prestin - molecular mechanisms underlying outer hair cell electromotility. Springer Handbook Of Auditory Research. New York: Springer.

Schanzler M, Fahlke C (2012) Anion transport by the cochlear motor protein prestin. J Physiol 590:259–272.

Sfondouris J, Rajagopalan L, Pereira FA, Brownell WE (2008) Membrane composition modulates prestin-associated charge movement. JBiolChem 283:22473–22481.

Song J, Santos-Sacchi J (2015) Intracellular calcium affects prestin’s voltage operating point indirectly via turgorinduced membrane tension. In: Mechanics of Hearing: Protein to Perception, pp 030009-030001. Cape Sounio, Greece: AIP Publishing.

Song L, Santos-Sacchi J (2010) Conformational state-dependent anion binding in prestin: evidence for allosteric modulation. Biophys J 98:371–376.

Song L, Santos-Sacchi J (2013) Disparities in voltage-sensor charge and electromotility imply slow chloride-driven state transitions in the solute carrier SLC26a5. Proc Natl Acad Sci U S A.

Tang J, Pecka JL, Tan X, Beisel KW, He DZ (2011) Engineered pendrin protein, an anion transporter and molecular motor. JBiolChem 286:31014–31021.

Tunstall MJ, Gale JE, Ashmore JF (1995) Action of salicylate on membrane capacitance of outer hair cells from the guinea-pig cochlea. J Physiol 485 (Pt 3):739–752.

Walter JD, Sawicka M, Dutzler R (2019) Cryo-EM structures and functional characterization of murine Slc26a9 reveal mechanism of uncoupled chloride transport. Elife 8.

Wang X, Yang S, Jia S, He DZ (2010) Prestin forms oligomer with four mechanically independent subunits. Brain Res 1333:28–35.

Yu X, Yang G, Yan C, Baylon JL, Jiang J, Fan H, Lu G, Hasegawa K, Okumura H, Wang T, Tajkhorshid E, Li S, Yan N (2017) Dimeric structure of the uracil:proton symporter UraA provides mechanistic insights into the SLC4/23/26 transporters. Cell Res 27:1020–1033.

Zhai F, Song L, Bai JP, Dai C, Navaratnam D, Santos-Sacchi J (2020) Maturation of Voltage-induced Shifts in SLC26a5 (Prestin) Operating Point during Trafficking and Membrane Insertion. Neuroscience 431:128–133.

Zheng J, Shen W, He DZ, Long KB, Madison LD, Dallos P (2000) Prestin is the motor protein of cochlear outer hair cells. Nature 405:149–155.

Zheng J, Du GG, Matsuda K, Orem A, Aguinaga S, Deak L, Navarrete E, Madison LD, Dallos P (2005) The C-terminus of prestin influences nonlinear capacitance and plasma membrane targeting. J Cell Sci 118:2987–2996.

Zivanov J, Nakane T, Forsberg BO, Kimanius D, Hagen WJ, Lindahl E, Scheres SH (2018) New tools for automated high-resolution cryo-EM structure determination in RELION-3. Elife 7.

